# CellPHIE: Integrating Pathway Discovery With Pooled Profiling of Perturbations Uncovers Pathways of Huntington’s Disease, Including Genetic Modifiers of Neuronal Development and Morphology

**DOI:** 10.1101/2025.02.19.639023

**Authors:** Byunguk Kang, Michael Murphy, Christopher W. Ng, Matthew Joseph Leventhal, Nhan Huynh, Egun Im, Serwah Danquah, David E. Housman, Ralda Nehme, Samouil L. Farhi, Ernest Fraenkel

**Affiliations:** Department of Biological Engineering, Massachusetts Institute of Technology, Cambridge, MA, USA; Spatial Technology Platform, Broad Institute of Harvard and MIT, Cambridge, MA USA; Computer Science and Artificial Intelligence Laboratory, Massachusetts Institute of Technology, Cambridge, MA, USA; Department of Biology, Massachusetts Institute of Technology, Cambridge, MA, USA; MIT Ph.D. Program in Computational and Systems Biology, Cambridge, MA, USA; Stanley Center for Psychiatric Research, Broad Institute of Harvard and MIT, Cambridge, MA, USA

## Abstract

Genomic screens and GWAS are powerful tools for identifying disease-modifying genes, but it is often challenging to understand the pathways by which these genes function. Here, we take an integrated approach that combines network analysis and an imaging-based pooled genetic perturbation study to examine modifiers of Huntington’s disease (HD). The computational analysis highlighted several genes in a subnetwork enriched for modifiers of neuronal development and morphology. To test the functional roles of these genes, we developed an experimental pipeline that allows pooled CRISPRi KD of 21 genes in human iPSC-derived neurons followed by optical analysis of genotypes, neuronal arborization, multiplexed pathway activity and morphological fingerprint readout. This approach recovered known genes involved in morphology and confirmed unexpected links from the network between several genetic modifiers of HD and morphology. Our approach overcomes challenges in pooled measurement of neuronal function and health and could be adapted for other phenotypes in HD and other neurological diseases.

## Introduction

Genome-wide association studies (GWAS) have identified many disease-modifying genes from high-throughput sequencing methods. The pathways by which GWAS hits function are challenging to understand from genomics alone. The combination of genomic screens and profiling methods, such as scRNA-seq and Cell Painting,^1–3^ have provided useful means to map between many genotypes and high dimensional phenotypes (cellular ‘fingerprints’). scRNA-seq fingerprints identify impacted transcriptional pathways well, but many functional phenotypes, particularly poorly annotated ones such as neuronal morphological changes are still challenging to understand. Cell Painting fingerprints, in contrast, provide evidence of phenotypic effects directly, but do not identify the molecular mechanisms by which these occur. Both methods can address these shortcomings in sufficiently scaled experiments which can contextualize results for perturbing genes of interest with similarity to the effects of perturbing other well understood genes.^4–7^ But progress has been much slower in phenotypes—such as those in neuronal diseases—which occur in cell types that are difficult to grow *in vitro* and involve complex morphological changes. Tackling this challenge in neuropsychiatric disease has become the focus of a consortium scale effort (SSPsyGene Consortium) but this is hard to replicate for every disease.^8,9^

We sought an approach that could be generally applied to neuronal diseases, focusing on Huntington’s disease (HD), which has not yet received the same level of attention. HD is an inherited and fatal neurological disease without a cure yet. It is caused by an inherited CAG repeat expansion in polyglutamine tract of the Huntingtin protein.^10^ Repeats longer than 40 give rise to disease and the length of CAG repeat tends to be inversely correlated with age of motor onset (AOO). However, this only explains ∼50% of the observed variability.^11–13^ After accounting for the CAG length, the “residual” AOO is a heritable trait.^14–16^ Multiple GWAS have identified potential AOO modifiers independent of the CAG length,^14–16^ but the mechanisms through which they influence HD pathophysiology remain unclear.

To nominate molecular mechanisms associated with HD AOO, we developed an integrated computational and experimental approach that can be conducted in medium-scale pools of dozens of gene knockdowns (KD) in induced pluripotent stem cell-derived neurons (iPSC-Ns). The current approach could be to perform genetic or morphological characterization after perturbing all AOO-associated genes, and then computationally finding commonalities from the data. We instead reasoned that the experimental load of pathway discovery can be decreased by first computationally nominating pathways through network analysis with subsequent experimental validation using a pooled genetic perturbation study (Figure 1). Thus, we present <u>Cell</u>-intrinsic <u>p</u>at<u>h</u>way discovery framework with <u>F</u>unctional, *In situ* profiling <u>E</u>xperiments (CellPHIE). First, we integrate knowledge from various studies by mapping genes onto a protein-protein and protein-metabolite interaction (PPMI) network, drawing on data generated across multiple model systems and data types (Figure 1A). This systematic mapping then allows us to identify relevant genes in a specific cell type and prioritize a functional phenotype to investigate by densely sampling genes from a subnetwork for experimental testing. We then developed an experimental toolbox based on multiplexed immunofluorescence (IF) to validate a key pathway by assessing both unbiased cellular fingerprints and targeted functional phenotypes (Figure 1B) in a pooled assay compatible with iPSC-Ns. We hypothesized that in this assay, interacting gene pairs should have similar fingerprints and should induce changes in the targeted functional phenotype associated with the overall network. We would thus be able to validate our computational results and establish relevance between the nominated genes and phenotype in a lower scale experiment.

**Figure 1.**
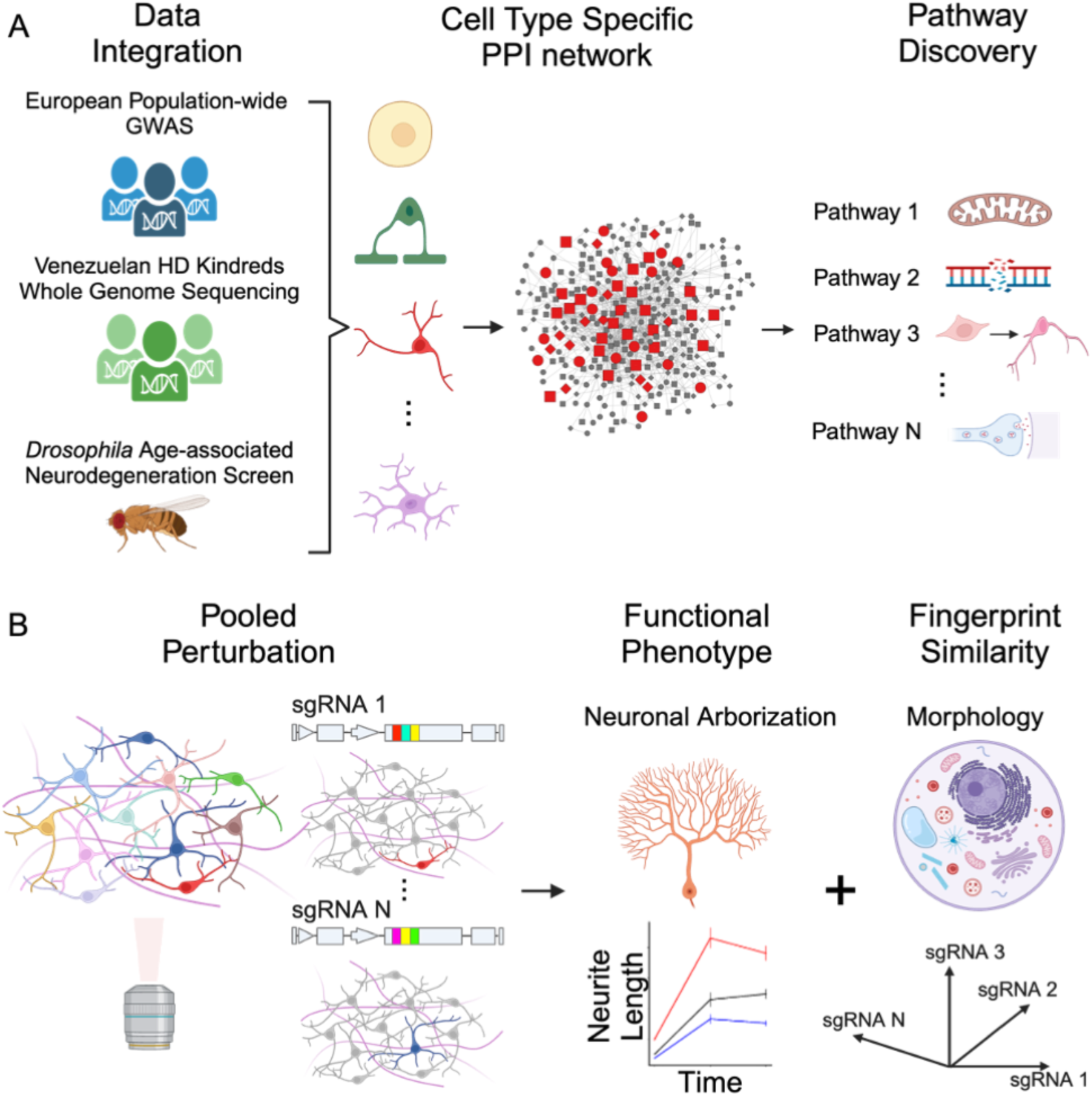
Overview of CellPHIE approach. A) Network modeling to integrate various human genetics datasets including GeM-HD GWAS data and Venezuelan HD Kindreds WGS study. To identify common mechanisms of neurodegeneration and HD, we also included modifiers of the rate of neurodegeneration (ND) onset in *Drosophila*. Genes were mapped onto a cell type specific PPI network and various pathways were identified. B) Pooled CRISPRi knockdown of genes densely sampled from a prioritized subnetwork, adapting an optical barcoding with Pro-Codes, followed by measurement of both unbiased morphological fingerprints and neuronal arborization phenotypes in iPSC-derived neurons.

In this study we demonstrate the CellPHIE framework by nominating putative AOO pathways in HD from a combination of various HD human genetics data and *Drosophila* neurodegenerative screens. Specifically, we included genetic modifiers of HD AOO identified from a population-wide, large scale GWAS data from the GeM-HD consortium^14^ and a whole genome sequencing data of a HD cohort from Venezuelan Kindreds accounting for rare variants.^17^ In addition to the HD data, we also included data from a genome-scale screen conducted in *Drosophila* that genes altering the rate of onset of neurodegenerative processes; we have previously shown that this screen revealed many drivers of human neurodegeneration (ND).^18^ Non-human model systems are particularly valuable in this context, as studies of human genetics may miss key modifiers because a variant may be too rare or toxic to appear in a human population. To systematically relate modifiers from various data sources and identify neuron-specific pathways of HD AOO, we adapted a previously developed, multi-omic network integration approach.^19^ Our network analysis revealed several neuron-specific pathways including neuronal development and arborization pathway, which we further validate experimentally.

To test the functional roles of the modifiers, we sought an unbiased readout of multiple cellular functions and a targeted readout of neuronal arborization pathways suggested by our network analysis. Since no single method could fulfill all these objectives, we integrated several methods into a unified experimental workflow. Our approach consists of pooled CRISPRi KD in iPSC-derived neurons, optical barcoding of each genotype with Pro-Codes,^20,21^ unbiased morphological fingerprint readout with Cell Painting, readout of specific pathways with multiplexed immunofluorescence (IF) and neurite outgrowth measurements. We used this rich dataset to confirm the gene involvement in neuronal development and arborization in several ways, based on both unbiased fingerprint analysis and targeted pathway specific measurements. Our work shows an integrative approach to discover disease-related pathways and provides an experimental platform to investigate their functional roles.

## Results

### Network integration of human genetic data and Drosophila screens identifies pathways that modify HD

The first step to applying CellPHIE to investigate HD AOO pathways was to compile known modifiers through an integrative analysis of various human genetics data. We started from a previous mega-analysis^17^ of combined genotypes and clinical data from the GeM-HD GWAS cohort^14^ primarily from a European population with many common and small effect size variants and the Venezuelan HD Kindreds whole-genome sequencing study, which consisted of rare variants with large effect sizes.^17^ We then systematically generated a list of AOO modifier genes identified from variants with functional consequence on protein function or gene expression (Table S1). We included genes in which SNPs significantly associated with altered AOO are likely to alter protein function (deleterious coding variants)^22–24^ or expression level (expression QTLs, eQTL) in prefrontal cortex utilizing the xQTL database.^25^ In total, we identified 218 HD AOO modifier genes. We recovered many confident modifiers previously identified from others, such as *FAN1*, *MLH1* and *MSH3*, all of which have been known to be involved in somatic repeat instability,^26–29^ one of the most widely studied pathways in HD.

We decided to supplement the human HD AOO modifiers with genes that affect the rate of onset of ND processes in general, and therefore may also play a role in HD relevant pathways. For this purpose, we leveraged a genome-scale genetic screen in *Drosophila*.^18^ This study identified 169 human-ortholog genes that accelerated ND, causing pathological changes to the brain after neuron-specific KD at an age of 30 days, approximately half of the fly life span.^30^ To relate modifiers of HD AOO and ND, we applied the Prize-Collecting Steiner Forest (PCSF) algorithm. PCSF is a multi-omic network integration approach that has been previously used to discover biological processes associated with many diseases.^18,19,31–35^ PCSF maps genes or molecules of interest from previous data onto a general human interactome of known or predicted protein-protein/-metabolite interactions (PPMI)^18,36–38^ and identifies part of the interactome most relevant to the input data, providing a comprehensive view of biological pathways spanning multiple data sources. This yielded a putative protein interaction network enriched for pathways that modify HD (Figure 2A).

**Figure 2.**
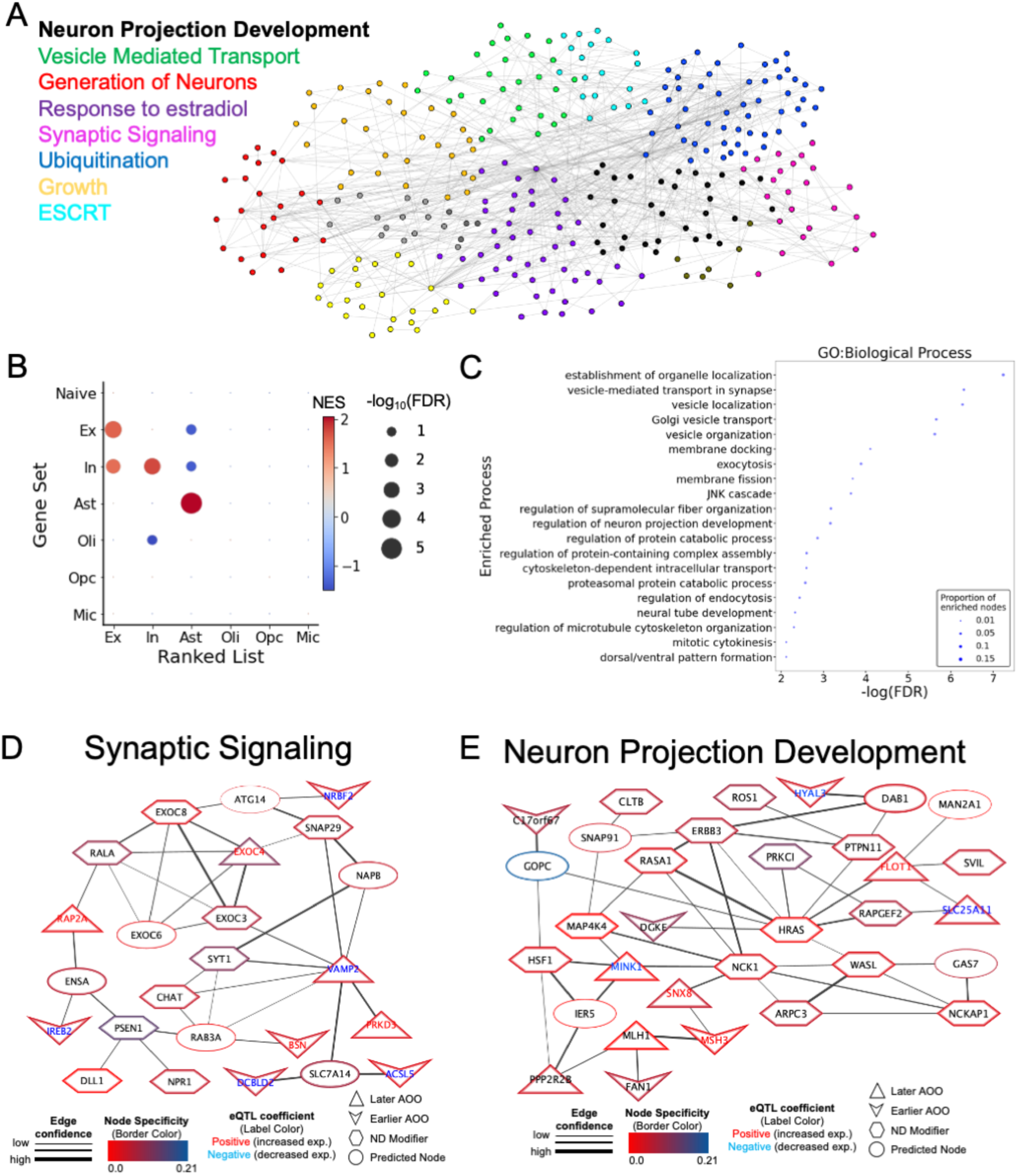
Network Integration of HD human genetics and forward genetic screen of altered rate of ND onset in *Drosophila* identifies putative pathways of HD. A) Overview of neuron-specific network output and enriched pathways. Nodes represent genes and edges represent interaction, with node colors matching the font colors of listed pathway. B) Gene set enrichment analysis (GSEA) of cell type specific gene expression profile demonstrating significant, positive normalized enrichment score (NES) of Ex, In, and Ast network nodes in their respective cell type specific DEGs. C) Overview of enriched Gene Ontology (GO) terms of neuron specific network and examples of subnetworks enriched for D) “synaptic signaling” and E) “neuron projection development” processes. For HD modifier genes (“Later AOO” and “Earlier AOO”), eQTL-associated genes are color-coded with node labels in red (increased expression) or blue (decreased expression) and genes identified from coding variants are labeled in black.

The PCSF output includes interactions across the entire proteome without regard to cell type. Since excitatory neurons (Ex) are known to be selectively vulnerable to HD pathology,^39^ we sought to further refine the list for Ex-specific pathways before experimental validation. We reasoned we could generate cell-type specific networks by adjusting the weights of each modifier and interactome edges based on their cell-type specific expression profile (Methods). To evaluate this approach, we ran PCSF separately for each of the major brain cell types including excitatory (Ex) and inhibitory (In) neurons, astrocytes (Ast), oligodendrocytes (Oli), oligodendrocyte progenitor cells (Opc) and microglia (Mic), utilizing previously published HD post-mortem brain single-nucleus RNA-seq (snRNA-seq) datasets.^17,40^ A cell-type naïve network was also generated without any weight adjustments, to assess cell type specificity. To evaluate cell type specificity of each network, we performed Gene-Set Enrichment Analysis (GSEA) (Figure 2B, Methods). Genes identified from Ex, In, and Ast networks showed significant, positive enrichment (FDR < 0.05) in differentially expressed gene lists (DEG) comparing gene expression profiles of each cell type to overall expression (Figure 2B, Methods), suggesting that these networks were indeed specific to their respective cell types. Importantly, the naïve network did not show any enrichment in any of the cell types used. We did not observe cell type specificity in other glial cell types (Oli, OPC, Mic), possibly because ND screen hits were identified from a neuron-specific RNAi KD screen^18^ and thus biased towards neuronal cell types. As expected, ND screen hits showed significant, positive enrichment (FDR < 0.1) in Ex neuron DEG list (Figure S1) while HD AOO modifiers were not enriched for any cell types. These results demonstrate the utility of our approach to identify cell type specific networks and to aid in targeted hypothesis generation.

Gene Ontology (GO) enrichment analysis of the genes identified in Ex network revealed several biological processes such as organelle localization, vesicle-mediated transport, JNK cascade, neuron projection development, proteasomal protein catabolic process and microtubule cytoskeleton organization (Figure 2C, Table S2, FDR < 0.05). The detailed results of this network are visualized in an interactive website (Data and Code availability). We also identified genes that were not initially included as modifiers of HD AOO or ND, but selected by PCSF (Predicted Node, Methods). These genes are likely relevant to the nominated pathways but may be missed by human genetics or *Drosophila* screen data, highlighting the discovery potential of our approach.

Previous research has shown that clustering of networks can often reveal functionally related groups of proteins.^18,41,42^ Thus, we applied Louvain community detection algorithm to PCSF output network to identify clusters of genes that are tightly interconnected with each other, and nominate possible pathways. In total, we identified 12 subnetworks and annotated their enriched pathways, summarized in Table S3. Many of the enriched pathways in these subnetworks were consistent with prior HD literature. For example, one subnetwork was enriched for processes related to synaptic signaling such as SNARE binding and vesicle-mediated transport in synapse (Figure 2D), in line with previous studies implicating synaptic dysfunction in HD.^43–45^ Importantly, significant enrichment for synaptic signaling was only observed after running PCSF, but not in the input set of genes (Figure 2C, Table S2), suggesting that our network analysis groups together genes involved in important biological processes.

One nominated subnetwork was of particular interest, as it was enriched for neuron projection development and actin cytoskeleton organization process (Figure 2E). Several studies have reported impaired actin cytoskeleton functions in the presence of mutant HTT (mHTT) or with loss of normal HTT,^46–49^ and others have found developmental defects resulting from these perturbations.^50–53^ Nonetheless, the impact of genetic modifiers on neuronal development in HD has been largely unexplored, especially compared to DNA damage and synaptic mechanisms. Thus, we sought to design an experimental system to test the functional roles of the modifiers in this hypothesized subnetwork in the neuron projection development pathway.

### Multiplexed immunofluorescence enables pooled profiling of neuronal morphology in iPSC-derived neurons

Evaluating the hypothesized network above would require performing perturbations of each nominated gene in a human system (e.g. iPSC-Ns); provide a range of well understood phenotypic readouts, including neuronal arborization; and be generally extendable to other future subnetworks. Because of the technical variability often observed in arrayed imaging assays, the ideal method would also allow all perturbations to be carried out in a pooled format to minimize experimental overhead and variability.^54–58^ No single technology can accomplish all these goals. Thus, we combined several approaches in a unified experimental pipeline consisting of perturbation with CRISPRi KD in iPSC-Ns; optical barcoding of each genotype with Pro-Codes; Cell Painting readout of unbiased morphological fingerprints; and targeted monitoring of specific pathways with multiplexed IF.

To perform pooled CRISPRi KD, we adapted an optical barcoding approach from Perturb-map^20,59^ and performed multiplexed IF to simultaneously identify the targeted gene in each cell and measure its neuronal morphology. We chose KD rather than knock out because of previous reports of knockout-associated toxicity in iPSC-Ns.^54,60–63^ Each targeted gene is labeled with a unique, triplet combination of epitopes (Pro-Codes) fused to dNGFR, a truncated transmembrane receptor without intracellular domain, which localizes to the membrane.^20^ This allows identification of both the cellular phenotype and the perturbed with iterated cycles of antibody staining (Figure 3A).

**Figure 3.**
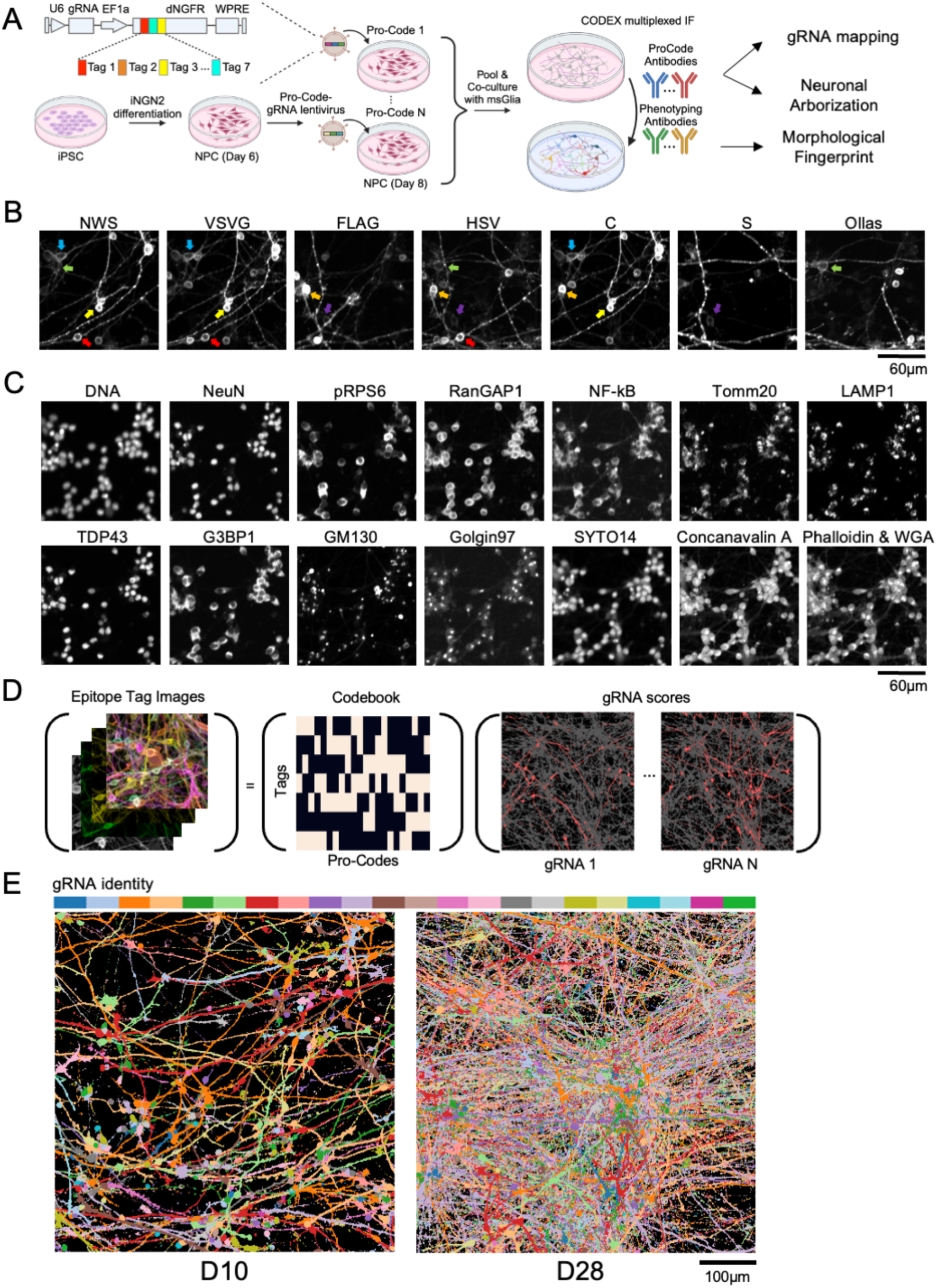
Pro-Codes enable *in situ* mapping of genotypes and phenotypes in iPSC-Ns. A) Overview of experimental workflow. B) Example field of view containing seven epitope tag images, where colored arrows highlight examples of cells stained positive with exactly three epitope tags. C) Same field of view as panel B showing additional phenotyping channels, combining antibody-based and Cell Paint dye-based measurements. D) Schematics of Pro-Code deconvolution problem. Matching Pursuit algorithm deconvolves raw tag images into a set of gRNA score images, given a codebook dictating triplet combinations of tag denoting each Pro-Code combinations. Each channel of the deconvolved image output represents the membrane label of each genotyped cell (colored in red), overlaid to cells of all other genotypes (colored in gray). E) Example field of view images (D10, D28) showing merged outputs of Matching Pursuit deconvolution algorithm of each genotyped cells assigned with 22 different pseudo-colors.

We first assessed whether the Pro-Code approach could be applied in iPSC-Ns to label gRNA identity and validated CRISPRi activity. We differentiated an iPSC line with a healthy donor background (WTC11) harboring doxycycline-inducible Neurogenin 2 (iNGN2) and dCas9-KRAB at the safe harbor locus^54^ into neurons using NGN2 overexpression and small molecule patterning towards a forebrain fate.^64,65^ We then transduced neurons with gRNAs targeting progranulin (GRN) or non-targeting control guides and observed robust knockdown of GRN via immunofluorescence (Figure S2).

Having confirmed that we can detect the effects of knockdown using Pro-Code vectors and an imaging assay, we individually cloned 44 gRNA-Pro-Code constructs targeting 21 genes and non-target controls (2 guides per gene, both assigned to the same Pro-Code, Figure 3A), packaged them into lentivirus, and transduced into day 6 (D6) early post mitotic neurons. 48 hours post transduction, cells were dissociated and pooled in equal ratio onto a monolayer of mouse glia and co-cultured up to D28 of neuronal differentiation (Figure 3A, Methods). We then performed CO-detection by inDEXing (CODEX),^66^ a commercial multiplexed IF technique to stain cells with 7 epitope specific antibodies to label the 22 Pro-Codes corresponding to each gene target (Figure 3B). Qualitatively, our results demonstrate that cells were indeed labeled with specific, triplet combination of epitope tags (Figure 3B, arrows).

To generate phenotypic information for these gRNAs, we sought to combine two types of measurements: unbiased fingerprinting and phenotype-specific measurements. For unbiased fingerprinting, we were inspired by the Cell Painting assay’s ability to generate morphological fingerprints predictive of common drug and gene mechanisms of action^2,67,68^. Thus, we chose to supplement the Pro-Code antibodies in the CODEX panel with both the organelle specific Cell Painting dyes (Hoescht, SYTO14, Concanavalin A, Phalloidin and WGA) and antibodies specific to additional organelles like the nuclear envelope (RanGAP1), lysosome (LAMP1), cis and trans-golgi (GM130, Golgin-97) and mitochondria (Tomm20) (Figure 3C). We further sought to add targeted phenotype-specific measurements which could directly test the phenotypes associated with the network. We reasoned that the membrane targeted Pro-Codes (Figure 3B) would permit eventual reconstruction of neuronal arborization phenotype associated with the nominated network, but we also added a set of more specific markers of HD relevant pathways such as active translation and mTORC1 pathway (pRPS6), neuronal maturation (NeuN), stress granules (G3BP1), NF-kB signaling (NF-kB), and RNA binding and metabolism (TDP43) (Figure 3C). The full CODEX panel consisted of 7 epitope channels, 10 phenotyping antibodies and 5 dyes.

To allow accurate mapping between gene perturbation and phenotype in densely packed neuronal cultures, we developed a novel de-barcoding method based on the Matching Pursuit sparse approximation algorithm (Methods). This allowed deconvolution of epitope tag images into a set of pixel-wise scores of each gRNA identity (Figure 3D, S3), even in mature neuronal cultures with densely packed neurites (Figure 3E). Next, we identified single-soma regions and assigned their corresponding genotypes based on maximum channel intensity of the deconvolved Pro-Code images (Figure S3, S4A). We then filtered out false assignments caused by neighboring cells or passing neurites (Figure S4B) using a logistic regression classifier that achieved high accuracy (AUC = 0.9, Figure S4C, Methods). 22 unique cell populations were identified, each characterized by a specific Pro-Code with expression of exactly three tags out of seven tags (Figure S4D,E). We recovered a total of 23,221 cells with clear Pro-Code assignment and associated morphological information. Taken together, our use of Pro-Code labeling combined with highly multiplexed IF readout enables labeling of genetically perturbed cells and measuring their morphology in a pooled culture.

### Morphological fingerprints capture functional similarity of HD modifiers

As Cell Painting is an assay that responds to a broad range of cellular functions, it provides an opportunity to explore functional similarities and differences among the genetic hits.^2,3,72^ For each identified cell’s single-soma region, we quantified various image features implemented in the Cell Profiler package^73^ including intensity, texture, correlation, granularity of each image channel and geometric features of cell segmentation masks (Figure 4A, S3, Method). We then took two approaches to investigate morphological features of the modifiers that emerged from our network analysis.

**Figure 4.**
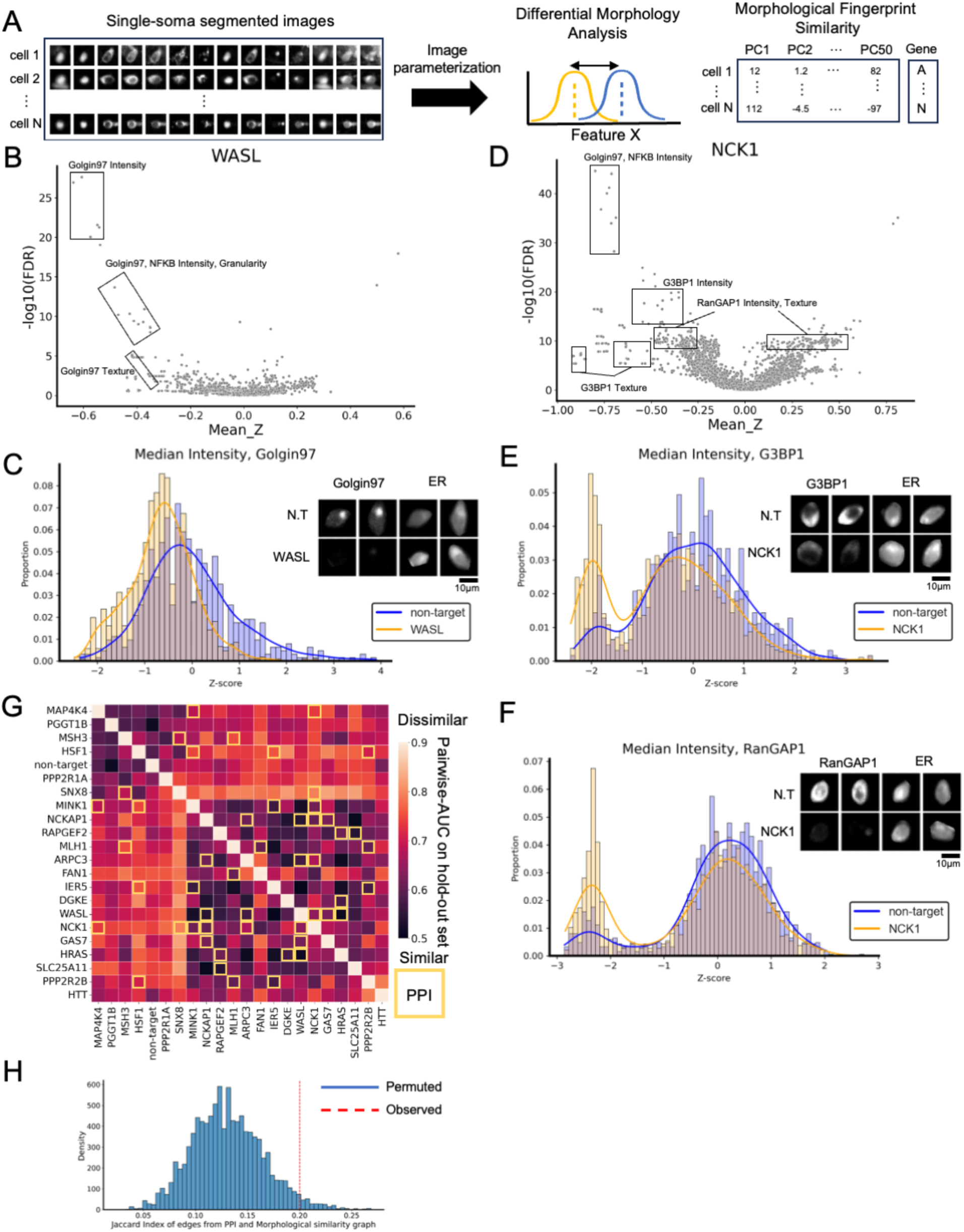
Multiplexed imaging panel identifies co-functional genes. A) Overview of single-soma image feature extraction and analysis workflow. B) Volcano plot of features comparing *WASL* KDs to non-targets (N.T). C) Histogram of the median intensity of Golgin97 channel comparing cells with guides targeting *WASL* and N.T. D) Volcano plot of features comparing *NCK1* KDs to N.T. E,F) Histogram of the median intensity of G3BP1 and RanGAP1 channel comparing cells with guides targeting *NCK1* and N.T. G) Pairwise gene-gene classification result. Row and column labels denote perturbed genes and each element represents AUC metric of classifier trained to classify between the two gene labels. Rows and columns are hierarchically clustered based on Pearson correlation. Gene pairs with known PPI are highlighted in yellow boxes. H) Jaccard Index between underlying PPI graph edges and measured similarity profiles (“observed”, red vertical line). The null distribution of permuted jaccard index is shown in blue histograms (“permuted”) where gene labels are permuted 10,000 times.

First, we asked which genes had strong effects on individual readouts. We performed differential morphology analysis, conceptually similar to differential gene expression analysis in transcriptomics studies.^72^ We conducted Kolmogorov-Smirnov (KS) test to assess whether there is a significant distribution shift between KD cells for each gene and non-target cells, for each feature individually (Method). We found that intensity-related features of Golgin97, NF-kB, RanGAP1 and G3BP1 were among the most shifted features in many perturbed genes. Reduced expression of both Goglin97 and NF-kB were shared in most of KDs (14 genes) while decreases in RanGAP1 and G3BP1 were shared in a subset of genes (8 genes) (Table S4). As an example, reduced Golgin97 intensity, was among the most differential features of *WASL* KD compared to non-target cells (Figure 4B,C), matching *WASL*’s known role and localization in trans-Golgi networks (TGN).^74^ KDs of *NCK1*, *SNX8*, *FAN1* and *PPP2R2B* gave the largest reductions of G3BP1 and RanGAP1 intensity (Figure 4D-F, S5). These results suggest changes in stress granules and nuclear-cytoplasmic transport, respectively. Both processes have previously been shown to be affected in HD models. Reduced levels of G3BP1 have been previously reported to induce mHTT aggregation in HD iPSC-Ns and *C. elegans* HD models.^75^ Furthermore, RanGAP1 levels were reduced in R6/2 HD mouse models.^76^ Our data suggest testable hypotheses for a deeper understanding of the mechanism of action underlying these phenotypes by highlighting genes which may mediate these effects.^75,76^

The analysis of individual features, particularly intensity-related features of each channel revealed several genes that share similar changes. However, the multidimensional output of our assay also allows us to map the pathways that are affected by HD genetic hits without a need to assume a reductionist one-gene-one-pathway approach. To explore such multi-pathway modifiers, we applied ‘morphological fingerprint analysis.’ These fingerprints are global representations of morphological cell state in response to each gene KD across all channels used in our assay. We examined the principal components of all Cell Profiler features for each cell to eliminate the influence of redundant, covarying features that could lead to spurious correlations. We then asked whether our multiplexed CODEX panels provide more complete morphological fingerprints, compared with those generated from conventional cell painting dyes alone. To that end, we assessed the extent to which the CODEX data help to distinguish between cells with non-target guides and each gene KD (Methods). Compared with using Cell Painting dyes alone, the use of multiplexed CODEX panels improved overall discriminative capabilities of our assay from mean AUC of 0.58 to 0.69 (Figure S6), averaged across all gene-targets. In some cases, the added benefit is much more dramatic. For example, the standard Cell Painting assay is of little use in distinguishing *SLC25A11* KD from non-target cells (AUC = 0.54), but the addition of the CODEX panel increases the AUC to 0.74 (Figure S6). Our results demonstrate that morphological fingerprints can indeed detect subtle responses to genetic perturbation.

Next, we assessed morphological fingerprint similarity between each gene KD pair using single-cell level classification performance (AUC) of logistic regression classifiers trained to discriminate between each gene labels (Figure 4G). The similarity profiles of each gene pairs were consistent across two replicate samples (Figure S7A), suggesting that our results were robust and reproducible. Importantly, morphological similarity was not correlated with similarity of epitope tag image intensities used to assign gene labels to each cell (Figure S7B), suggesting that our results reflect biological similarity rather than technical artefacts.

We observed several groups of genes that show similar morphological profiles (AUC < 0.65, Method) while being distinct from non-target controls (Figure 4G). These groups of genes are thus likely to be involved in the same pathway, functioning similarly. We found similar profiles between *MLH1* and *FAN1* KD (AUC = 0.60), consistent with their known interaction to promote resistance to DNA interstrand cross-link damage and preventing formation of cytotoxic DNA cleavage products.^77^ We also found similar profiles between *IER5* and *PPP2R2B* KD (AUC = 0.57), consistent with their known mutual interaction with protein phosphatase 2A (PP2A) to alter downstream target phosphorylation.^78–82^ Our results recovered not only similarity profiles of known physical interaction partners, but also those of functionally relevant gene pairs. We found strong similarity profiles of *IER5* and *MLH1* KD (AUC=0.52), both of which have been previously shown to sensitize cells to radiation-induced double-strand DNA breaks.^83,84^ On the other hand, we observed modestly distinct profiles between *IER5* and *FAN1* (AUC = 0.68), consistent with *IER5*’s association with non-homologous DNA end joining (NHEJ) repair pathway^83^ versus *FAN1*’s association with DNA repair by homologous recombination (HR).^85^ Importantly, while we did not directly measure DNA damage or mismatch repair activities in our assay, morphological fingerprinting approaches such as Cell Painting has been previously reported to be predictive of DNA damage phenotypes and other features related to cell health,^67^ supporting our assay’s capability to capture functionally relevant relationships as well as physical interactions.

Finally, to evaluate whether our morphological fingerprints recovered expected PPI relationships, we quantified the amount of overlap between PPI edges from the nominated subnetwork (Figure 2E) and edges derived from morphological fingerprint similarity graph (Jaccard index, Methods). The morphological similarity profiles indeed recovered many known interactor edges from the PPI network model, with a Jaccard index of 0.2 (empirical p-value = 0.03, Figure 4H), suggesting that our nomination method can faithfully recover biological interactions. Taken together, these results support our assay’s ability to recover expected relationships from PPI networks. More importantly, the overall process so far allows us to start from a computationally nominated list of genes likely to share a common pathway and find experimental evidence to support that. For example, we find morphological similarities between *SLC25A11* and *NCKAP1*, suggesting that these genes are associated with an HD-related neurite outgrowth pathway, despite a lack of previous literature examining these genes in this context.

### Pooled neuronal arborization analysis recapitulates known regulators and identifies novel regulators of neurite outgrowth

We next set out to test our hypothesis that the genes in the subnetwork would alter neuronal development and arborization when knocked down (Figure 2E). Rather than the traditional neuronal arborization analysis that involves sparse, arrayed transfection with fluorescent proteins^54^ or staining neurites with axonal and dendritic markers,^70,71^ we reasoned that we could use the membrane localized Pro-Code signals to recover the same information in a pooled format (Figure 3E). Thus, we repurposed deconvolved Pro-Code images and adapted a previously developed, automated neurite tracing pipeline^54^ to measure neuronal arborization phenotypes such as neurite length, number of branches and number of trunks (Figure 5A) for each KD. Here, each channel of deconvolved Pro-Code images corresponds to each genotype and can be processed separately (highlighted in red, Figure 3D), distinguished from other neurons of unrelated genotypes (highlighted in gray, Figure 3D). This labeling approach provided several advantages over traditional, arrayed neuronal arborization assay. First, our approach reduced experimental load by pooling multiple genotypes into a single culture. Second, sparsely labeling individual neurons allowed robust tracing of each neurite without being obscured by the neighboring neurites. As expected, we observed neurite outgrowth during the first 21 days of differentiation, with fewer changes at later times (p-value < 1e-12, two-tailed t-test comparing neurite length of non-target cells in D10 and D28, Figure 5B). The effects of each gene KD were also highly consistent across replicate samples across two independent experiments (Figure S8), suggesting that our pipeline was robust in accurately recovering neuronal arborization parameters.

**Figure 5.**
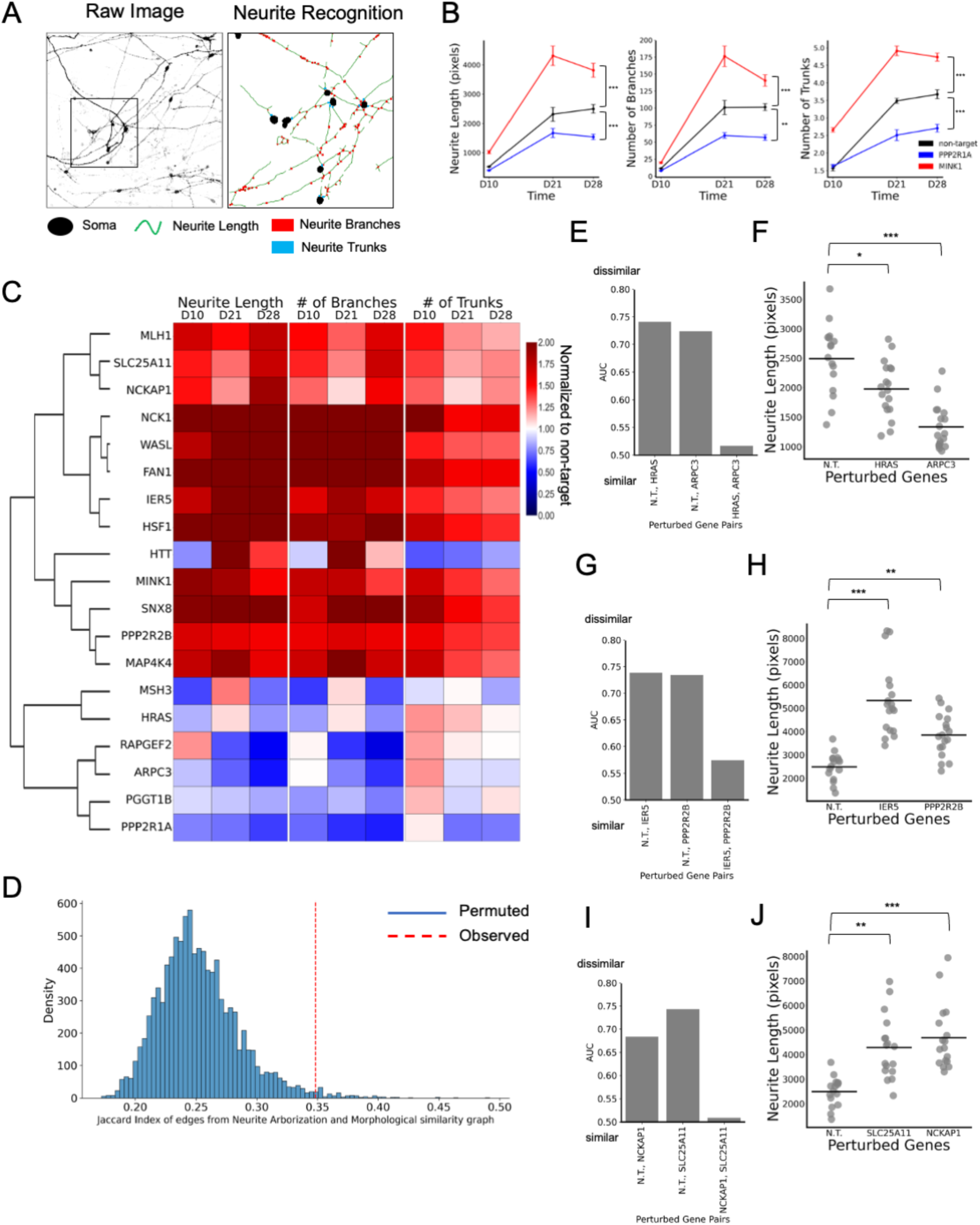
Pooled neuronal arborization analysis recapitulates known regulators and identifies novel regulators of neurite outgrowth. A) Deconvolved Pro-Code images can be used to extract neuronal arborization features such as neurite length, number of branches, and number of trunks, demonstrated by (left) single-channel example corresponding to the orange channel in Figure 3E (D10) and (right) zoomed-in view of neurite feature recognition. B) Effects of knockdown of MINK1 (positive regulator) and PPP2R1A (negative regulator) on neurite outgrowth across different timepoints. C) Effects of knockdown of different genes (row) on neuronal arborization features (neurite length, number of branches, and number of trunks) relative to non-targeting gRNAs at different time points (columns). Rows were hierarchically clustered based on Pearson correlation. D) Jaccard index between edges derived from soma fingerprint similarity graph and arborization similarity graph (“observed”, red vertical line). The null distribution of permuted jaccard index is shown in blue histograms (“permuted”) where gene labels are permuted 10,000 times in soma fingerprint similarity graph. E-J) AUC similarity metric of soma fingerprint features for each gene pair (E, G, I) and neurite length profiles (F, H,J). Each dot is a technical replicate and horizontal line represents mean value. For panels B,F,H and J, two-tailed t-tests were performed and significant differences compared to non-targeting gRNAs are reported with p-value corrected for multiple hypothesis testing, denoted as *** FDR < 0.00001, ** FDR < 0.001 and * FDR< 0.01.

Consistent with our initial hypothesis from network analysis that the genes that we profiled are involved in neuronal development and neurite outgrowth pathways, we observed significantly altered neuronal arborization patterns in many gene knockdowns in this subnetwork when compared to non-target control. These effects broadly clustered into positive and negative modifiers of neurite outgrowth phenotypes (Figure 5C). Among the negative modifiers, we highlight that *PPP2R1A* knockdown resulted in the reduction of neurite length and the number of branches and trunks (D28, FDR < 1e-6 for all parameters using two-tailed t-test, Figure 5B, S9), consistent with a previous study in the same WTC11 line.^54^ We did not observe the increased neurite length in *PGGT1B* KD in the same study but instead observed increased number of trunks at a comparable timepoint (D10, FDR<0.05). This discrepancy could be due to differences in KD timing, differentiation protocol, and culture conditions. We also recovered literature results among the positive modifiers, such as finding that knocking down *MINK1* and *MAP4K4* resulted in increased arborization (D28, FDR < 1e-3 for all parameters using two-tailed t-test, Figure 5B,C, S9), consistent with previous studies employing genetic or pharmacological inhibitions in various neuronal model systems.^86–89^

We then asked whether Cell Painting, a broad measure of cellular state (morphological fingerprint), might pick up signals in the soma that are correlated with arborization. To assess the consistency between the two assays, we quantified the amount of overlap between gene pairs that resulted in consistent arborization phenotypes and those derived from morphological fingerprint similarity graph from previous section (Jaccard Index, Methods). The similarity profiles observed in soma features indeed were consistent with similarity in neuronal arborization phenotypes, with a Jaccard index of 0.35 (empirical p-value = 0.01, Figure 5D). These results, therefore, would suggest that morphological fingerprint indeed captures functional relationships observed in an orthogonal direct phenotyping assay.

As an example of the consistency between somatic and neurite measurements, we highlight that *HRAS* and *ARPC3* KD shared highly similar morphological fingerprints extracted from single soma multiplexed CODEX images (AUC = 0.52), while each gene KD effect was highly distinguishable from non-target controls (Figure 5E). Consistent with their known functions related to neurite outgrowth^90–94^ and their morphological similarity observed in soma images, *HRAS* and *ARPC3* KD resulted in significant reduction of neurite length and number of branches compared to non-target controls (Figure 5F). Our result would thus provide further evidence that the two genes are involved in a shared pathway.

Going further, *PPP2R2B* and *IER5* KDs shared highly similar morphological fingerprints extracted from single-soma multiplexed CODEX images (AUC = 0.57, Figure 5G), resulted in increased neuronal arborization phenotype (Figure 5H). The effect of *PPP2R2B* KD on arborization was consistent with a previous study that demonstrated reduction of neurites upon overexpression of *PPP2R2B*.^95^ This result is also consistent with transcriptomic measurement in a Perturb-seq study previously performed in the same WTC11 line, where positive regulation of gene modules involved in “axon guidance” pathway was observed.^58^ In contrast, little is known about *IER5*’s involvement in neuronal arborization pathway. Our results from two orthogonal assays suggest that *IER5* and *PPP2R2B* share functional similarity and *IER5* may indeed affect neuronal arborization.

Finally, our results also highlight similar perturbation responses among genes that are not predicted to interact directly or do not belong to previously known pathways, but may inform us of candidate targets for further investigation in the context of HD. For example, the morphological fingerprint of *NCKAP1* KD was indistinguishable from that of *SLC25A11* (AUC = 0.51, Figure 5I), even though these two genes are not predicted to interact and their functional relevance to each other is not known previously. Reduced expression of *SLC25A11* in human patients was associated with later age of onset (eQTL associated gene, Figure 2E), suggesting a putative protective effect of reduction of *SLC25A11* in HD. Our data would, therefore, suggest that *NCKAP1* expression would modify HD. Indeed, *NCKAP1* KD in a mouse model of HD enhanced neuronal survival.^96^ Interestingly, both genes showed an increased neuronal arborization phenotype (Figure 5J), providing additional evidence that the two genes may be related. Taken together, these results suggest that morphological fingerprint indeed captures functional relationships and is useful even in cases where it is difficult to assay the phenotype directly.

## Discussion

In this study, we demonstrate that CellPHIE can identify disease pathways by integrating computational nomination of pathways and experimental validation of a key pathway through unbiased morphological fingerprinting and targeted functional readouts.

We first bring together insights from multiple datasets across different model systems and generate hypotheses on cell-type specific pathways and relevant functional phenotypes to measure in a less resource-intensive approach compared to other experimental efforts. We used PCSF network analysis to integrate hundreds of modifiers of HD AOO that emerged from various human genetics datasets and modifiers of neurodegeneration emerged from *Drosophila* screens, and systematically identified several pathways of HD. Our integrative analysis helped complement human genetics data that may miss important modifiers on its own and identify converging, common pathways of general neurodegenerative process and HD-specific process. Future investigations may include additional evidence from other model systems, such as a screen for modifiers of mHTT-mediated toxicity in mouse models, to further understand mechanisms of actions of these modifiers.^96^ We further refined our discovery process by generating cell type specific networks, ensuring that the nodes (proteins) and edges (interactions) are relevant to the cell type of interest and thus reducing false discoveries. Finally, we narrowed our focus by identifying a subnetwork enriched for neuronal development and neurite outgrowth pathway based on GO analysis.

We used two approaches to define phenotypes that demonstrate how nodes within the same subnetwork were related to each other. First, we tested whether we observe similar response in an unbiased, generic readout of cellular functions. Second, we sought to directly measure a specific, relevant functional readout such as neuronal arborization phenotype. To address these questions simultaneously, we developed an experimental pipeline that combines various profiling approaches – *in situ* genotyping with Pro-Code labeling in a pooled CRISPRi assay in iPSC-Ns, unbiased morphological fingerprinting with Cell Painting, readout of specific pathways with multiplexed IF and a measure of neuronal arborization. As expected, most of the genes—but not all—showed a similar unbiased morphological fingerprint. Furthermore, the genes showing this similar morphological fingerprint also had a common arborization phenotype, as predicted from the network nomination, as predicted.

Our network approach facilitates generating targeted hypotheses and pinpointing specific, functional measurements for experimental validation at multiple stages. For example, differential morphological analysis identified several genes in which key processes affected in HD models are impacted upon KD, such as changes in proteins involved in nuclear-cytoplasmic transport or stress granules. Our unbiased choice of multiplexed antibody panels also enabled us to identify processes that are relatively unknown in the process of ND or HD pathophysiology, such as changes in Golgi functions, to be investigated further in future studies. Moreover, morphological fingerprint analysis revealed functional similarities among HD modifiers that were consistent with their known functional relationships or interactions, demonstrating the utility of our multiplexed antibody panels. As an example, we observed similar morphological profiles between *IER5* and *MLH1* KD. *MLH1* is a highly confident HD modifier that is known to be involved in somatic repeat instability,^27,97,98^ while little is known for *IER5*, a predicted node that emerged from our network analysis and has not been shown previously to be implicated in HD. Through ‘guilt-by-association’ with the *bona fide* modifier of HD AOO such as *MLH1*, *IER5* may be an interesting target for further investigation.

Neuronal arborization analysis confirmed the effects of known modifiers of neurite outgrowth and validated the predicted involvement of the HD modifiers in neuronal arborization pathway, suggested by the network analysis. Notably, paired measurement of morphological fingerprints in soma regions and functional measurement of neuronal arborization yielded several novel hypotheses to be investigated in future studies. For example, our morphological guilt-by-association analysis revealed similar profiles between *SLC25A11* and *NCKAP1*, two genes previously not known to share a mechanism. Interestingly, they also shared similar neuronal arborization patterns, providing additional evidence that they may be functionally related. Thus, our data may explain their potential mechanisms of actions towards their beneficial impact on HD phenotypes - later AOO in human genetics (*SLC25A11*) and enhanced survival in HD mouse models (*NCKAP1*).^96^ Furthermore, *FAN1*, a high-confident HD modifier known to be involved in somatic repeat instability^26,27,29^ and *NCK1*, a ND modifier of yet unknown relationship to HD pathophysiology, resulted in the strongest change in soma and arborization phenotypes (Table S4). Little is known about *FAN1*’s relevance to neuronal outgrowth pathway. However, *FAN1* is also a risk gene for Autism Spectrum Disorder (ASD),^99–102^ a neurodevelopmental disorder, and the arborization phenotype that we observe is consistent with *FAN1* having a role in developmental processes. Future work will be needed to determine if these processes are also relevant to HD.

Our study has implications for the growing field of morphological screening. Morphological profiling, such as Cell Painting, has provided useful readout of cellular health and function, but its application in neurodegenerative disease has remained challenging, due to difficulties in growing relevant cell types *in vitro* and their complex morphological changes. Neuronal arborization parameters have been considered important indicators of neuronal health and function,^70,103,104^ but previous studies have been limited to arrayed characterizations with low throughput and relied on use of axonal or cytosolic markers for neurite tracing.^54,71^ Our use of Pro-Code membrane labeling not only identified the perturbed cells but also quantified neuronal arborization parameters in a pooled CRISPRi assay, providing the first description of a pooled neuronal arborization assay. We also, show, however, that high dimensional, generic morphological fingerprinting can perform similarly well in discriminating between genes as these arborization measurements. This suggests that, if the purpose of a study is to identify perturbations sharing a morphological pathway, a more technically tractable somatic based assay can be sufficient. This opens the door to high content optical pooled screening based screens based on neuronal morphology, as well as applications of the CellPHIE approach to larger numbers of subnetworks simultaneously in the same experiment.^3,72,105,106^

While our assay has provided useful information on cell health and functional phenotypes, it also has two important limitations. First, while our emphasis was on moderate-throughput screens, we recovered more cells than necessary for adequate power, showing that the number of profiled genes could be increased. The Perturb-map vectors can readily accommodate >300 genes, provided they are recloned to include a selection marker to remove the large number of unmodified cells. This would allow pooled transduction as well as pooled screening, and facilitate profiling of multiple subnetworks simultaneously. Second, we carried out the experiments in a wild type (WT) line, without mHTT and the negative effect of aging. We believe that uncovering the function of these modifiers in WT lines is a crucial first step to understanding them in the context of disease. Future studies could use isogenic lines expressing different lengths of CAG repeats to understand in greater detail how these genes interact with HD processes. Even so, our work focuses on identifying pathways through which AOO risk genes act and enriches for these genes from baseline morphological profiling data—suggesting these pathways are active in WT cells.

In sum, this study establishes a framework to computationally nominate disease relevant pathways and experimentally validate key pathways through high-dimensional image-based profiling methods coupled with neuronal arborization measurements. Through iterative network modelling and experimentation, we believe that our approach will serve as a valuable platform and resource to study disease pathways.

## Methods

### Data availability

The full network is available and explorable at: https://github.com/bkang-mit/CellFIE/tree/master/subnetwork_html

### Code availability

Code can be found at https://github.com/bkang-mit/CellFIE

### Cell type specific network integration

To identify pathways through which modifiers of HD AOO and ND contribute to HD pathophysiology, we applied the Prize-Collecting Steiner Forest algorithm (PCSF) implemented in OmicsIntegrator 2 package (version 2.4.0).^19^ Briefly, the algorithm maps modifier genes to a reference ‘interactome’ graph, a set of known protein-protein and protein-metabolite interactions (PPMI) derived from iRefIndex version 17, HMDB, Recon 2 and BraInMap databases as previously described.^18,36–38^ This mapping is controlled by ‘prize’ - user-defined weights reflecting disease-relevance of the molecules of interest and ‘edge cost’ – reflecting the degree of support for the PPMI. By maximizing prizes and minimizing edge costs, the algorithm identifies the part of the interactome most relevant to the input datasets. As these reference interactions were defined in human proteins and metabolites, the genetic hits derived from *Drosophila* dataset were mapped to human orthologs as previously reported.^18^ We assigned prizes to HD modifiers based on their significance and effect sizes. ND modifiers were assigned with median prize value of HD modifiers.

The output of PCSF, as implemented in OmicsIntegrator 2 package (version 2.4.0) is a cell type naïve network and may include molecules and their interactions that are irrelevant to the cell type of interest. We reasoned that adjusting the weights of prizes and cost of interactome edges based on cell type specific gene expression profiles can generate more targeted hypotheses relevant to the cell type of interest. For each cell type, previously published HD post-mortem brain snRNA-seq datasets^17,40^ were utilized to calculate enrichment score of prizes (*E*) and specificity score of interactome edges (*S*) (Equation 1). Enrichment score (*E*) reflects the relative expression level of a given prize *j* in a cell type *t* compared with overall expression levels in the tissue, and specificity score (S) between gene *i* and *j* is a penalty term applied to the edge cost determined by the rank percentile of a gene expression level within the cell type *t*. To preserve prize weights based on prior knowledge (e.g. relative impact on HD AOO) and the likelihood of protein-protein interaction, these scores were scaled with fixed weights and added onto cell type naïve prize and edge cost (Equation 2).

**Equation 1**. Cell type enrichment score of prizes (E) and specificity score of edges (S), where 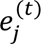 is expression level of gene *j* in cell type *t* and 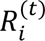 is rank percentile of gene *i* in cell type *t*.

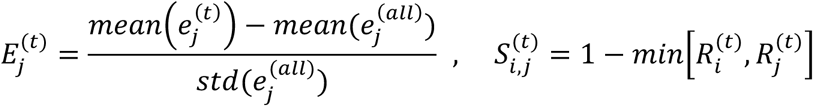

**Equation 2**. Cell type specific prize (P) and edge costs (C) with hyperparameter K.

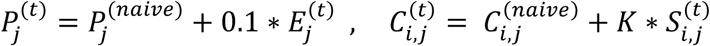

We ran PCSF separately for each cell type prize and edge cost adjustments. For each network, we performed 100 randomizations of the edges with gaussian noise, to assess robustness of the nodes. We also performed 100 randomizations of prize values to assess specificity of each node to their prize assignments. We then retained nodes that appeared more than 50 of the robustness randomizations and more than 50 of the specificity randomizations. To select optimal hyperparameters for OmicsIntegrator, we evaluated various ranges; β = {50, 100}, determines scaling factor applied to prizes, γ = {4,5,6}, determines the edge penalty regularizing highly connected ‘hub’ nodes, *K* = {0.05, 0.1, 0.15}, determines the relative contribution of cell type specific edge penalty. Final networks were chosen based on minimizing the mean specificity, maximizing the mean robustness, and minimizing the KS statistic between node degree of the prizes as compared to those of the predicted nodes, as previously described.^18^ Our final selection was β = 50, γ = 5, *K = 0.1*

Finally, to evaluate cell type specificity of each network, we performed gene set enrichment analysis (GSEA), assessing whether nodes are enriched for genes that are highly expressed in each cell type. Subnetworks were then identified through Louvain community detection clustering (resolution = 1) of the resulting, larger PCSF selected network for pathway analysis. Gene ontology (GO) term enrichment was performed using gProfiler package (version 1.0).

### Molecular cloning and lentivirus packaging

Each gRNA oligo was cloned individually into corresponding backbone Pro-Code Vector (Table S5, Gift of Brian Brown) as previously described.^20^ To produce lentivirus, 1e+6 HEK293T cells were plated in each of 6-well tissue culture plates with DMEM (Gibco, Cat. No. 11965092) supplemented with 10% FBS. On the next day, cells were transfected with 1.6 μg of gRNA-Pro-Code construct, 0.4 μg of pMD2.G plasmid (VSV-G envelope, Addgene #12259), and 1 μg of psPAX2 plasmid (Addgene #12260), using TransIT-293 transfection reagents (Mirus Bio, Cat. No. MIR2704) according to the manufacturer’s protocol. Media was changed to Opti-MEM media (Gibco, Cat. No. 11965092) at 18hr post transfection and viral supernatant was harvested at 48 hr post transfection. All guide sequences and their matching Pro-Code constructs are listed in Table S5.

### NGN2 neuronal differentiation and pooled co-culture

Human iPSCs engineered to express mNGN2 under a doxycycline-inducible system in the AAVS1 safe harbor locus and dCas9-KRAB in the CLYBL intragenic safe harbor locus^54^ (male WTC11 background, gift from Michael Ward) were used in this study. Neuronal differentiation was carried out following previously established protocols with some modifications.^64^ On Day 1, iPSCs were seeded at a density of 40,000 cells/cm² in tissue culture plates coated with Geltrex (Gibco, Cat. No. A1413301) using mTESR medium supplemented with ROCK inhibitor and 2 µg/ml Doxycycline hyclate (Sigma, Cat. No. D9891-25G). On Day 2, the medium was replaced with Neuronal Induction media (NIM), which included DMEM/F12 (Gibco, Cat. No 11320-033), N2 supplement (Life Technologies, Cat. No. 17502048), 20% Dextrose (VWR, Cat. No. BDH9230-500G), Glutamax (Life Technologies, Cat. No. 35050079), Normocin (Invivogen, Cat. No. Ant-nr-2), 100 nM LDN-193189 (Stemcell Technologies, Cat. No. 72147), 10 µM SB431542 (Stemcell Technologies, Cat. No. 72234), 2 µM XAV (Stemcell Technologies, Cat. No. 72674), and 2 µg/ml Doxycycline hyclate. NIM was refreshed on Day 3. On Day 4, the medium was switched to Neurobasal Media (NBM, Life Technologies, Cat. No. 21103049) containing B27 supplement (Gibco, Cat. No. 17504044), MEM Non-Essential Amino Acids (Life Technologies, Cat. No. 11140076), Glutamax, 20% Dextrose, 2 µg/ml Doxycycline hyclate, Normocin, 10 ng/ml BDNF (R&D Systems, Cat. No. 11166-BD), 10 ng/ml CNTF (R&D Systems, Cat. No. 257-NT/CF), and 10 ng/ml GDNF (R&D Systems, Cat. No. 212-GD/CF). The cells were cultured for an additional 2 days before lentivirus transduction.

At Day 6, cells were transduced individually with lentivirus expressing each gRNA-Pro-Code pairs. The following day, medium containing lentivirus was replaced with fresh medium. At Day 8, cells were dissociated with Accutase (STEMCELL Technologies, Cat. No. 7920), pooled at equal ratio and plated total of 1e+6 cells onto a mono-layer of mouse primary glia (p3) cultured in glass coverslips (Akoya Biosciences, Cat. No. 7000005) coated with 0.1% Poly-L-Lysine (Sigma-Aldrich, Cat. No. P8920) and Geltrex. Medium was changed every 3-4 days, with the same NBM formulations as above up to Day 28.

### Quantification of knockdown by immunofluorescence imaging

To assess dCas9-KRAB activity and knockdown effects, we individually transduced Day 6 early post mitotic neurons with lentiviral constructs expressing gRNA targeting progranulin (GRN) and non-targeting control guides. Medium was changed to fresh medium the following day and cultured up to Day 10. Cells were fixed with 4% PFA in NBM media for 30 minutes at room temperature. Cells were then washed three times with 1x PBS, permeabilized with 0.1% TritonX in 1X PBS for 10 minutes at room temperature and blocked with 4% BSA in 1x PBS for 1 hour at room temperature. Cells were incubated in primary antibody cocktail containing anti-VSVg, anit-FLAG and anti-GRN antibodies (Table S5, 1:1000 dilution) in 4% BSA in PBS overnight at 4°C. On the next day, cells were washed three times with 1x PBS and incubated with secondary antibody cocktail containing DAPI nuclear counterstain (Invitrogen, Cat. No. D3571), Donkey anti-rabbit Alexa Plus 647 (Invitrogen, Cat. No. A32795), Donkey anti-rat Alexa Plus 647 (Invitrogen, Cat. No. A48272), and Donkey anti-goat Alexa Plus 555 (Invitrogen, Cat. No. A32816) at room temperature for 1 hour. Cells were then washed with 1x PBS for three times and imaged using an Andor Dragonfly 200 Spinning Disk Confocal Microscope with a Plan Apochromat 20x/0.75 NA objective (Nikon, MRD00205).

### PhenoCycler/CODEX Assay and imaging

At each timepoint, cells were fixed and washed as above and stained with CODEX barcode conjugated, primary antibody cocktails (Table S5) according to the manufacturer’s instructions. Antibody catalogs and conjugated fluorophores are listed in Table S5. After staining, each coverslip was imaged using Andor Dragonfly 200 Spinning Disk Confocal Microscope with a Plan Apochromat 20x/0.75 NA objective (Nikon, MRD00205), over multiple cycles of reporter incubation, imaging and stripping utilizing the CODEX Instrument Manager software and custom Python triggering script. 10 z-stacks spanning 10 μm across a large tile area (225 fields-of-view images, approximately 9 mm x 9 mm) were acquired. After 8 cycles of CODEX imaging rounds, samples were stained with Hoescht 33342 (Invitrogen, Cat. No. H3570), Concanavalin A (Invitrogen, Cat. No. C21401), SYTO14 (Invitrogen, Cat. No. S7576), phalloidin (Invitrogen, Cat. No. A12380) and Wheat-germ agglutinin (WGA) (Invitrogen, Cat. No. W32464), as previously described^2^ and imaged. A typical imaging run lasted 15-20 hours for each sample.

### Image preprocessing

Raw images were maximum-intensity-projected along z-dimension and registered using pystackreg package (version 0.2.5). Single-soma and nuclei were segmented using CellPose model^107^ (version 3.0) with custom human-in-the-loop training, using SYTO14 channel and DAPI channel. 14-channel phenotyping images were used to extract high-dimensional image features using custom CellProfiler (version 4.2.1) pipeline, broadly consisting of features related to area, shape, intensity, texture, granularity, colocalization between two channels and subcellular localization (radial distribution). For Pro-Code deconvolution, z-stack images were used to improve deconvolution results. 7-channel tag images were converted into [0, 1] range using robust quantile normalization using quantile ranges [75%, 99.9%] and Matching Pursuit algorithm (see below) was applied to generate 22-channel deconvoluted images, for genotyping and neuronal arborization feature extraction. Preprocessing workflow is summarized in Figure S3.

### Pro-Code deconvolution

We sought to deconvolve epitope tag images (*M*, 7-channels) into Pro-Code images (*S*, 22-channels, where pixel values of each channel correspond to membrane label intensity of each genotyped cell, given a codebook (*A*, 22 Pro-Codes by 7 tags) that dictates specific, triplet combinations of tags into Pro-Codes (Figure 3D). In other words, we aimed to find S that minimizes objective function: ‖𝐴,𝑆 − 𝑀‖^-^, the squared L2-norm distance between observed tag images *M* and approximated, projected image *A^T^S*. To solve this objective function, we adapted ‘matching pursuit’ sparse approximation algorithm as summarized below.

**Algorithm 1**. Pseudocode for Matching Pursuit Sparse Approximation of Pro-Code Images

*Input*:

- *M*: multi-channel image of dimensions (*m, z, y, x*) where *m* is number of measurement epitope channels and *z,y,x* are dimensions of image pixels.
- *A*: codebook matrix of dimensions (*n, m*), where *n* is number of output Pro-Code channels (genotypes).
- *max_iters*: maximum number of iterations, set to 3
- *scale_factor*: tunable parameter for optimal mask generation, set to 0.25

*Output*:

- · *S:* deconvolved Pro-Code image of size (*n, z, y, x*)

*Algorithm:*

1. Normalize each row of *A* to unit norm
2. Initialize S ← zero matrix of size (n, z, y, x)
3. Compute initial pixel-wise L2 norm: 𝑀_𝑛𝑜𝑟𝑚 ← ‖𝑀‖_-_ across channels
4. Compute foreground threshold: 𝑡ℎ𝑟𝑒𝑠ℎ𝑜𝑙𝑑 ← Otsu_threshold( 𝑀_𝑛𝑜𝑟𝑚) × 𝑠𝑐𝑎𝑙𝑒_𝑓𝑎𝑐𝑡𝑜𝑟
5. Compute foreground pixels: 𝑎𝑐𝑡𝑖𝑣𝑒_𝑠𝑒𝑡[𝑝] ← 𝑡𝑟𝑢𝑒 𝑓𝑜𝑟 𝑎𝑙𝑙 𝑝𝑖𝑥𝑒𝑙𝑠 𝑝 𝑤ℎ𝑒𝑟𝑒 𝑀_𝑛𝑜𝑟𝑚 > 𝑡ℎ𝑟𝑒𝑠ℎ𝑜𝑙𝑑
6. Initialize residual to input image: 𝑟 ← 𝑀
7. **for** t = 1 to *max_iters* **do**:

1. Project active pixels onto dictionary:ℎ ← 𝐴 ∗ 𝑟[: , 𝑎𝑐𝑡𝑖𝑣𝑒_𝑠𝑒𝑡]
2. Find maximum projection per pixel: 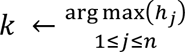
3. Update Pro-Code image: 𝑆[𝑘, 𝑎𝑐𝑡𝑖𝑣𝑒_𝑠𝑒𝑡] ← ℎ[𝑘]
4. Update residual: 𝑟[:, 𝑎𝑐𝑡𝑖𝑣𝑒_𝑠𝑒𝑡] ← 𝑟[:, 𝑎𝑐𝑡𝑖𝑣𝑒_𝑠𝑒𝑡] − 𝐴[𝑘], ∗ 𝑆[𝑘, 𝑎𝑐𝑡𝑖𝑣𝑒_𝑠𝑒𝑡]
5. Update pixels that converged: 𝑎𝑐𝑡𝑖𝑣𝑒_𝑠𝑒𝑡 ← {𝑡𝑟𝑢𝑒 𝑓𝑜𝑟 𝑝𝑖𝑥𝑒𝑙𝑠 𝑤ℎ𝑒𝑟𝑒 ‖𝑟‖_-_> 𝑡ℎ𝑟𝑒𝑠ℎ𝑜𝑙𝑑}

### Single-cell Barcode assignment and quality control

For each segmented single soma region, a genotype (Pro-Code) was assigned based on maximum channel intensity of deconvolved Pro-Code images. This intensity-based assignment can result in false barcode assignments due to neighboring cells, passing neurites, and other imaging artefacts summarized in Figure S4. To filter out these false assignments, approximately 2300 cells were manually labelled spanning across five different fields of view randomly sampled, assessing whether a barcode is assigned correctly or not based on maximum intensity channel of deconvolved images. Various metrics were quantified including overall tag intensity signal, per-soma area covered by Pro-Code reconstructed images, differences between third and fourth largest tag intensity, and the ratio of proportion of variance explained by top two principal components of 7-tag image intensities. Annotated data was then split into 70% training set and 30% test set and trained a logistic regression classifier (scikit-learn package, version 1.6) to classify each single-soma crops whether Pro-Codes are correctly assigned (Figure S4A) or incorrectly assigned due to artefacts (Figure S4B). Model’s performance was assessed in held-out cells (test set, Figure S4C). Cells that are correctly assigned are retained for further analysis.

### Differential Morphology and Morphological Fingerprint Similarity Analysis

The output of Cell Profiler pipeline consists of 23,221 cells by 2827 image features with Pro-Code (gene) mapping. For each replicate sample (coverslip), we computed the mean (µ) and standard deviation (σ) of feature *j*, within control cells and used these to z-normalize raw feature matrix *x*, such as 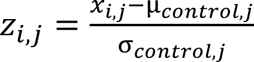 for *i*-th cell. To prevent the impact of outliers or numerical artifacts, we clipped the resulting z-scores to a range of [-10, 10] as commonly practiced in other high-dimensional, single-cell analysis.^108^ Two-sample Kolmogorov-Smirnov (KS) test was applied to determine whether there is a significant shift in distribution of each feature between KD cells for each gene and non-target cells as previously described.^72^ Features with clear trends in significant features were annotated.

To generate morphological fingerprint, principal component analysis was performed on the z-normalized cell-by-feature matrix, as many features were covarying and redundant. Top 50 principal components (PC) were retained to capture majority of variance and were defined as morphological fingerprints. To quantify similarity across each gene pair, data were subset to contain only the given gene labels and trained a binary logistic regression classifier to discriminate one gene label from the other. The data were split with 80% used for training and 20% held-out for testing. To address potential class imbalances in the data, the ‘balanced’ class weight setting was used in the ‘LogisticRegression’ function implemented in scikit-learn package (version 1.6), which adjusts the weights inversely proportional to class frequencies. The classifier’s performance was evaluated using area under the receiver operating characteristic curve (AUC-ROC), which can be interpreted as the degree to which two populations are distinct from each other, at a single-cell level. This metric typically ranges from [0.5, 1] where 0.5 indicates that the two populations are indistinguishable and 1 indicates that they are perfectly distinct. Values below 0.5 represent cases where the classifier performs worse than random chance and we did not observe any pair with AUC below 0.5.

To determine a threshold above which the two groups of cell populations can be considered to have distinct morphological fingerprints, non-target cell population was randomly assigned with two labels, shuffling their assignment 1000 times and each time a logistic regression classifier was trained to classify between the two randomly assigned labels. As expected, we observed mean AUC of 0.5 and the maximum AUC was 0.65. Thus, gene pairs with AUC less than 0.65 were considered to have similar morphological fingerprints. To calculate consistency between morphological fingerprint similarity and PPI network edges, a similarity graph was constructed by taking all gene pairs below AUC of 0.65 as edges. Jaccard Index (intersection divided by union) of the edges of similarity graph and PPI graph was calculated (‘observed’). To assess statistical significance of this consistency, the node labels of similarity graph were permuted 10000 times and Jaccard Index was calculated each time to construct a null distribution (‘permuted’). Empirical p-value was determined by calculating how many times ‘observed’ is greater than ‘permuted’ Jaccard index (Figure 4H).

### Quantification of neuronal arborization phenotype

To quantify arborization parameters of each genotype, Pro-Code deconvoluted images were binarized based on maximum channel (genes) intensity, closed with 5-pixel footprint and skeletonized (implemented in scikit-image package version 0.19.3). Skeletonized images were then masked out with soma segmented images and converted into network graph objects (implemented in networkx package version 3.4.2). Nodes that do not overlap with soma boundary are recognized as “branches” and those overlap with soma boundary are recognized as “trunks.” Total edge length is used as “neurite length” (Figure 5A). Small graph components of less than 5 nodes and 100 total edge pixel length are removed for robust quantification. Total of 25 field-of-view images were stitched and quantified and considered as one technical replicate among 9 others (total of 225 images per well replicate) and normalized to the total number of nuclei detected to account for cell density differences.

To quantify consistency across neuronal arborization assay results and soma morphological fingerprints, arborization graph was constructed by taking all gene pairs with consistent arborization direction. Jaccard index between edges derived from arborization graph and morphological fingerprint similarity graph was calculated as above. Null distribution and empirical p-value were calculated as above, by permuting morphological fingerprint similarity graph (Figure 5D).

### Statistics

Two sample, two-sided Kolmogorov-Smirnov test (KS-test) was applied for results in Figure 4 and Student’s t-test (two-tailed) was applied in Figure 5. For multiple hypothesis test corrected p-values, Benjamini-Hochberg FDR adjusted values were used.

## Author contributions

B.K., S.L.F., E.F. conceived of and designed the study. B.K. and M.M. developed algorithms for Pro-Code deconvolution method. C.W.N. and D.E.H. performed HD human genetics data analysis. B.K. and M.J.L. performed network integration and designed cell type specific adjustment methods. B.K. and S.D. designed and performed neuronal differentiation and imaging experiments with guidance and supervision from R.N. B.K. performed all data analysis, with assistance from N.H. and E.I.. B.K., S.L.F. and E.F. wrote the manuscript.

## Acknowledgements

We thank Michael Ward for providing the iPSC line used in this work. We thank Brian Brown for providing Pro-Code backbone vectors. This work was supported by NIH grants R01NS089076, R01MH128366 and Simons Foundation Autism Research Initiative award #890477 to SLF and RN.

## Supplementary Data

**Supplementary Table S1**. List of HD AOO modifiers with functional annotation used in this study (HD_AOO_Compiled_Annotation tab).

**Supplementary Table S2.** Gene Ontology annotation of PCSF output and input.

**Supplementary Table S3.** Enriched pathway of each subnetwork, including Gene Ontology, Reactome and KEGG pathway databases.

**Supplementary Table S4.** KS test result of intensity related features of each channel, sorted by p-value.

**Supplementary Table S5.** gRNA sequences, Pro-Code barcode assignment and catalogs for primary antibody and CODEX reagents used in this study.

## Supplementary Figure

**Figure S1.**
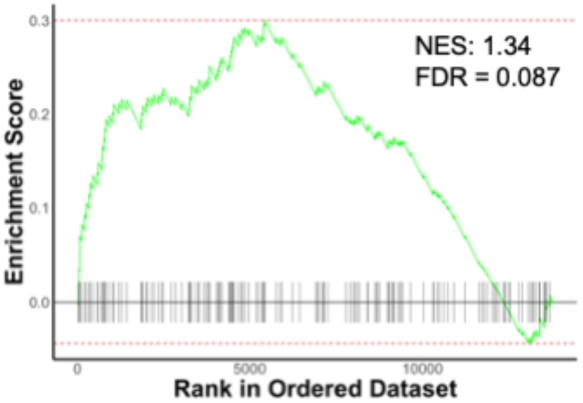
GSEA enrichment plot showing significant, positive enrichment of ND modifiers in excitatory neuron DEGs.

**Figure S2.**
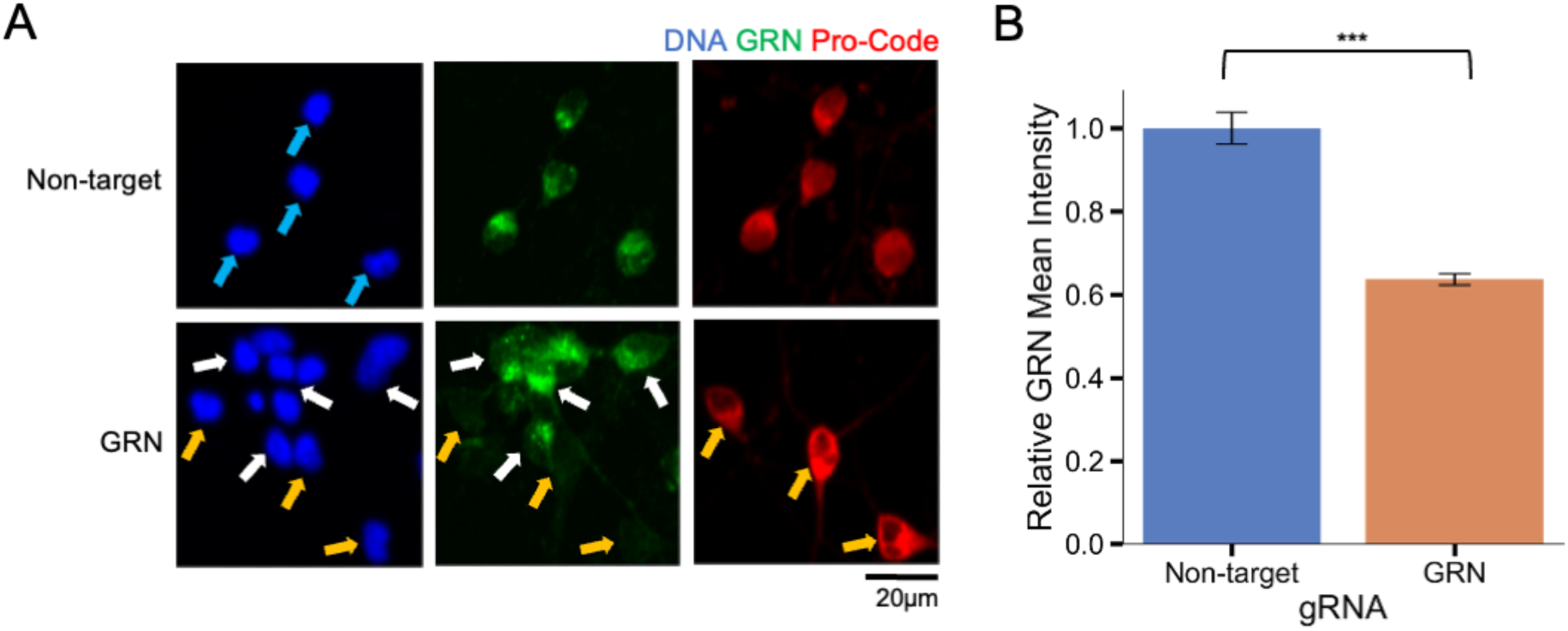
Validation of Pro-Code vectors and CRISPRi activity using immunofluorescence. A) Top row, non-targeting negative control gRNA. Bottom row, gRNA targeting progranulin (GRN). GRN signal (green), nuclear counterstain DAPI (blue), and Pro-Code tag (red) are shown. Blue and orange arrows highlight cells expressing gRNA-Pro-Code vector and white arrows highlight cells not expressing the vector. B) Quantification of GRN protein level, normalized to non-target controls. Bars and error bars represent mean ± S.E. (n=30 cells) (*** p-value < 1e-11, two tailed t-test)

**Figure S3.**
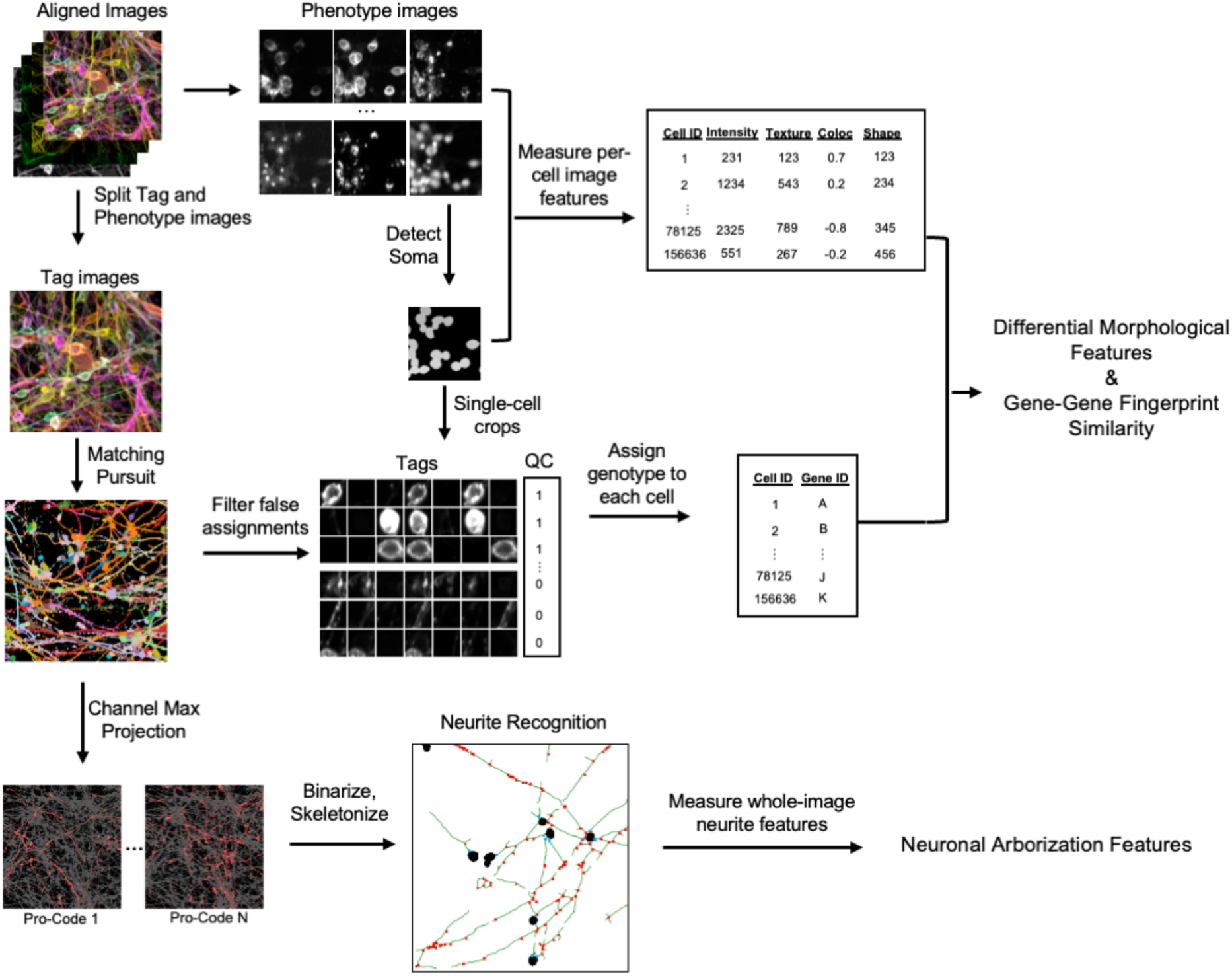
Overview of image processing workflow. Aligned images were split into tag and phenotyping images. Matching Pursuit algorithm was applied to deconvolve 7-channel tag images into 22-channel Pro-Code images where each channel represents each genotype used in this study. The deconvolved Pro-Code images were then used in each of segmented single soma regions to assign genotypes. Image features were extracted from 14-channel phenotyping images and combined with the single-soma genotype metadata to quantify single-cell morphological features. To quantify neuronal arborization parameters of each gene KD, each channel of Pro-Code deconvolved image is binarized and skeletonized, followed by custom neurite recognition pipeline (Method).

**Figure S4.**
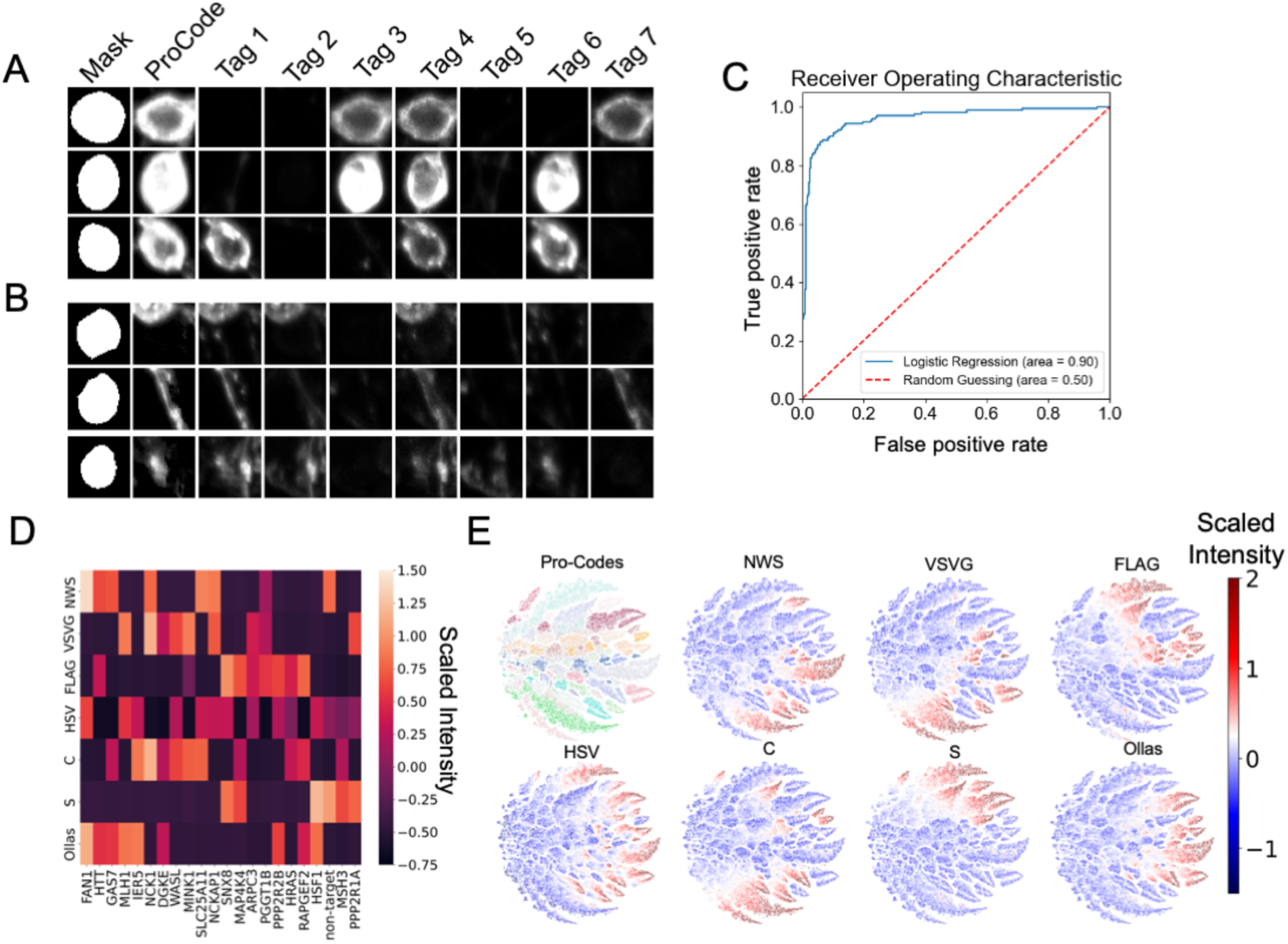
Post-processing of Pro-Code deconvolution for single-soma genotyping. Example cells that are A) correctly and B) incorrectly assigned with Pro-Code deconvolution images. Cells are falsely assigned with a given Pro-Code due to various imaging artifacts such as signals originating from neighboring cells or neurites and ambiguous signals from multiple tags. C) Receiver operating characteristic curve demonstrating performance of logistic regression classifier distinguishing correct assignments from false assignments. D) Scaled mean intensity of each of 22 genotyped single-soma regions. Rows denote each of epitope tags and columns denote gene names (Pro-Codes) corresponding to each triplet combination of epitope tags. E) TSNE plot demonstrating specific, triplet tag expression of each single cell.

**Figure S5.**
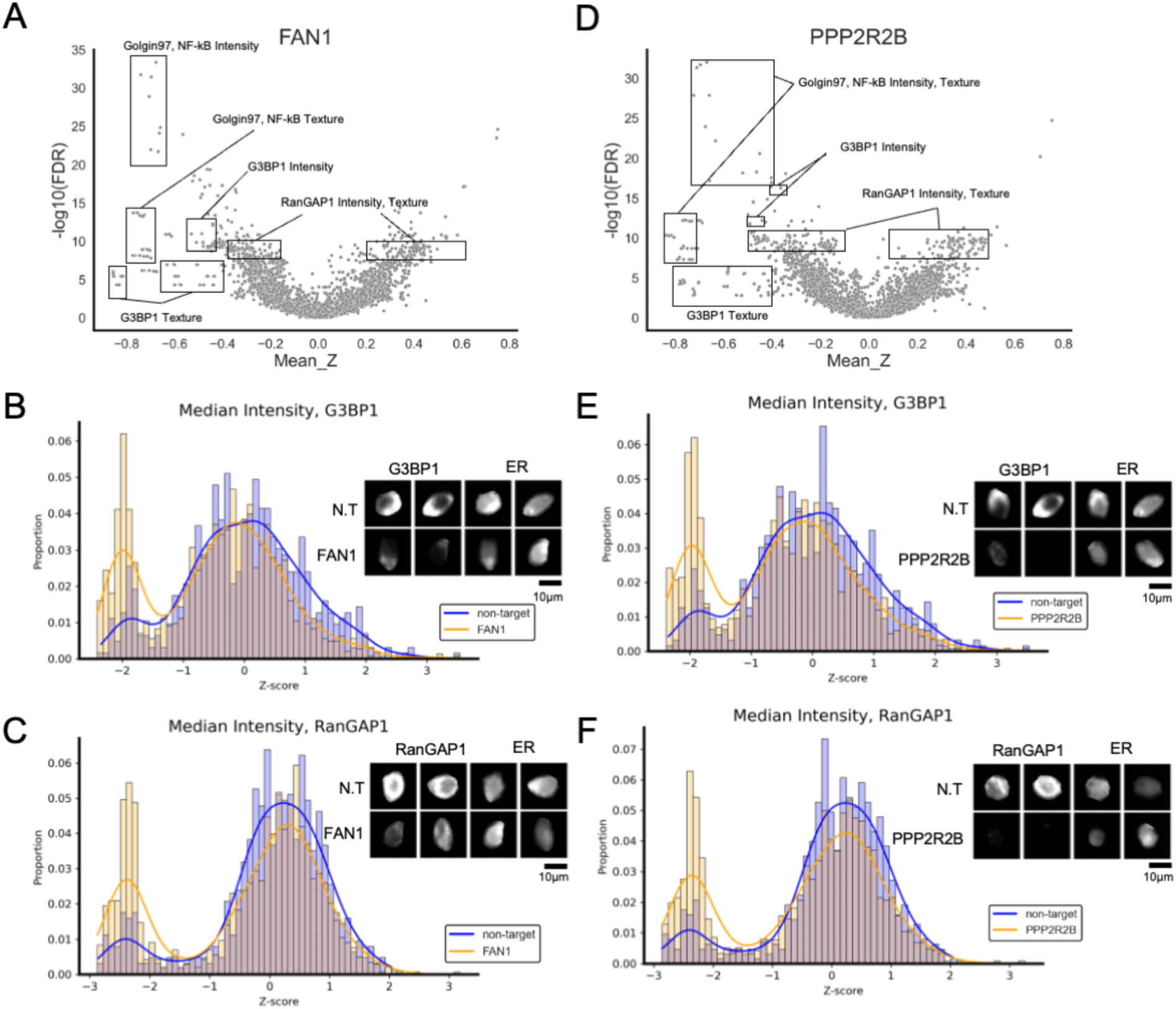
A) Volcano plot of features comparing *FAN1* KDs to non-targets (N.T). B-C) Histogram of the median intensity of G3BP1 and RanGAP1 channel comparing cells with guides targeting *FAN1* and N.T. D) Volcano plot of features comparing *PPP2R2B* KDs to non-targets. E-F) Histogram of the median intensity of G3BP1 and RanGAP1 channel comparing cells with guides targeting *PPP2R2B* and N.T.

**Figure S6.**
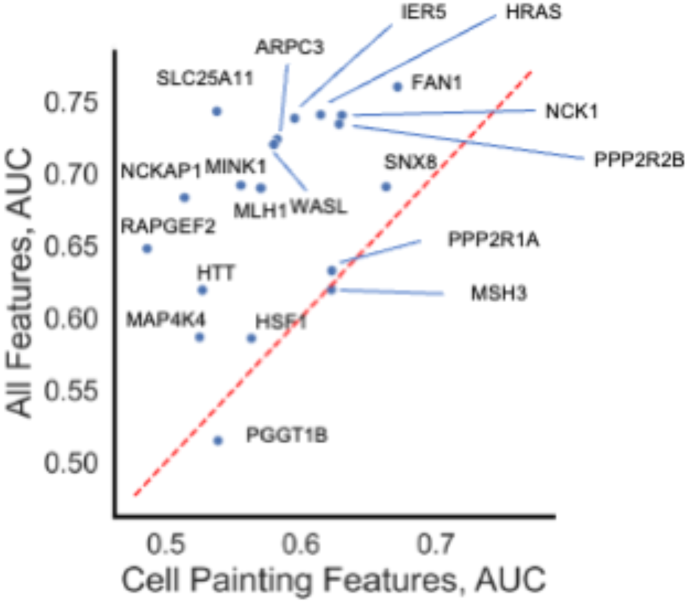
Comparison of features derived from Cell painting dyes only (x-axis) and all imaging panels including antibodies and dyes (y-axis), in their ability to distinguish between each gene KD (single dots) and non-target cells (AUC).

**Figure S7.**
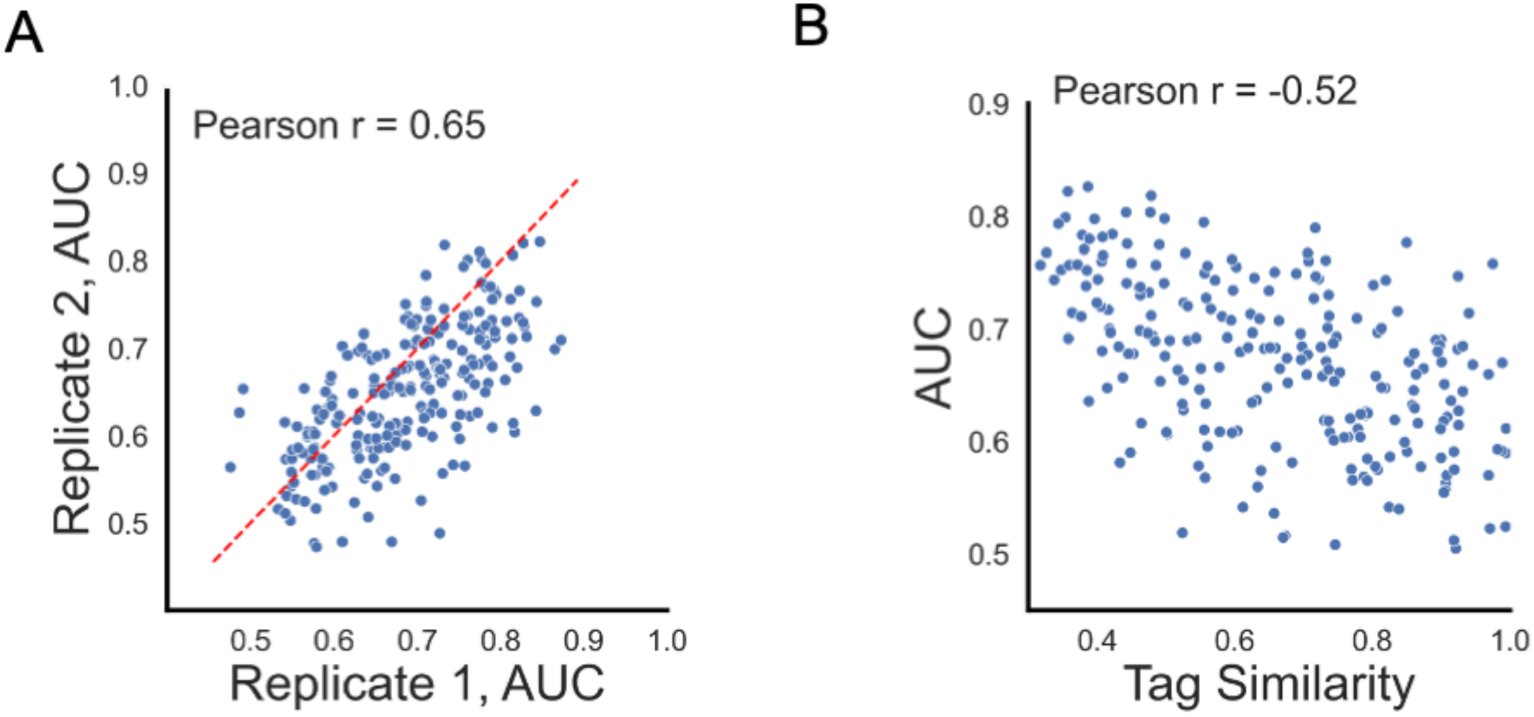
A) Replicate-replicate correlation of gene-gene pairwise AUC metric. B) Comparison of gene-gene similarity profiles using morphological fingerprint features (pairwise AUC metric, y-axis) and cosine similarity of epitope tag intensity (x-axis). Each dot represents each gene-gene pairs in panels A-B.

**Figure S8.**
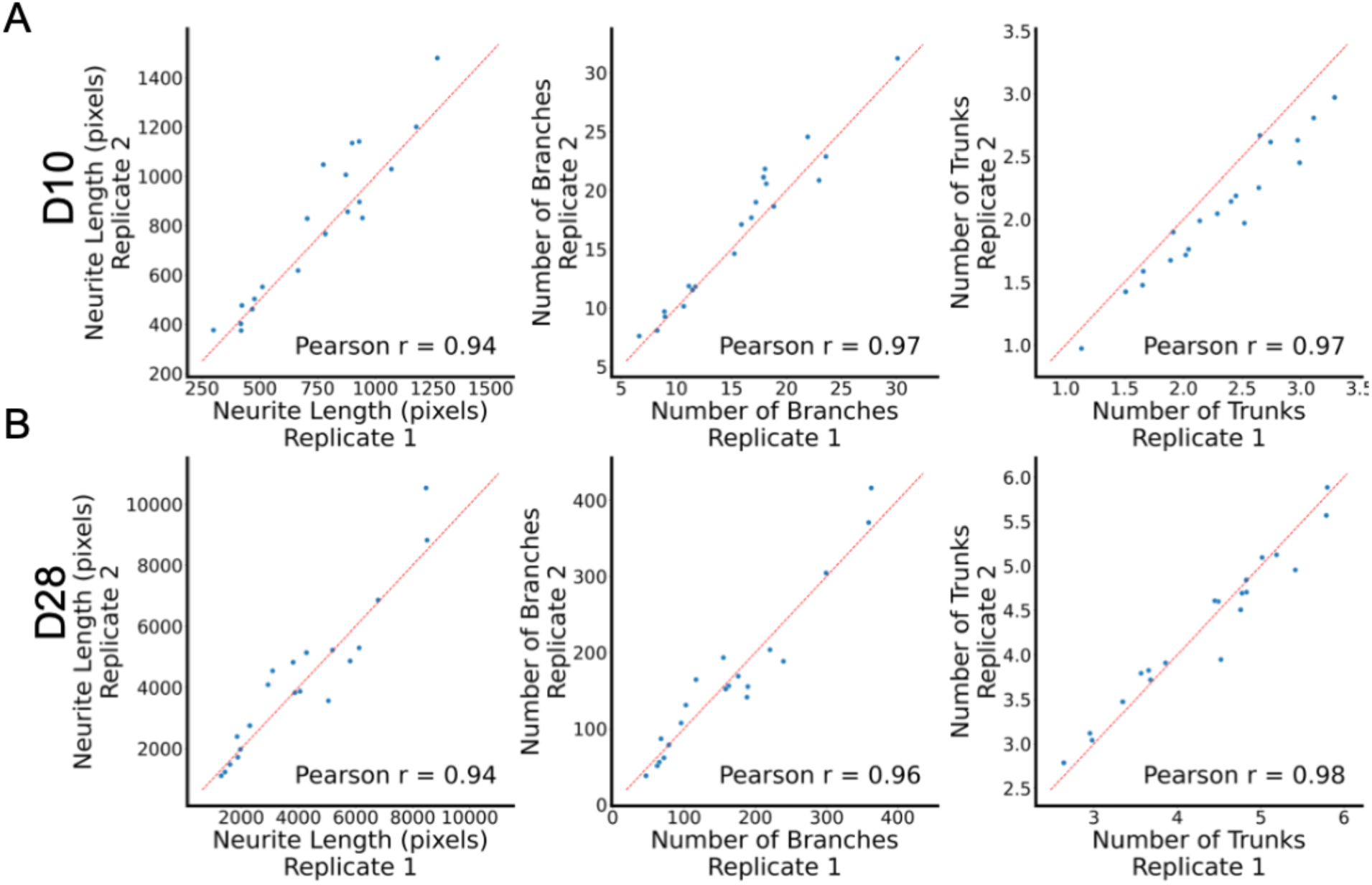
Replicate-replicate correlation of neuronal arborization features in A) Day 10 and B) Day 28 samples. Each dot corresponds to the mean feature values for each gene KD condition.

**Figure S9.**
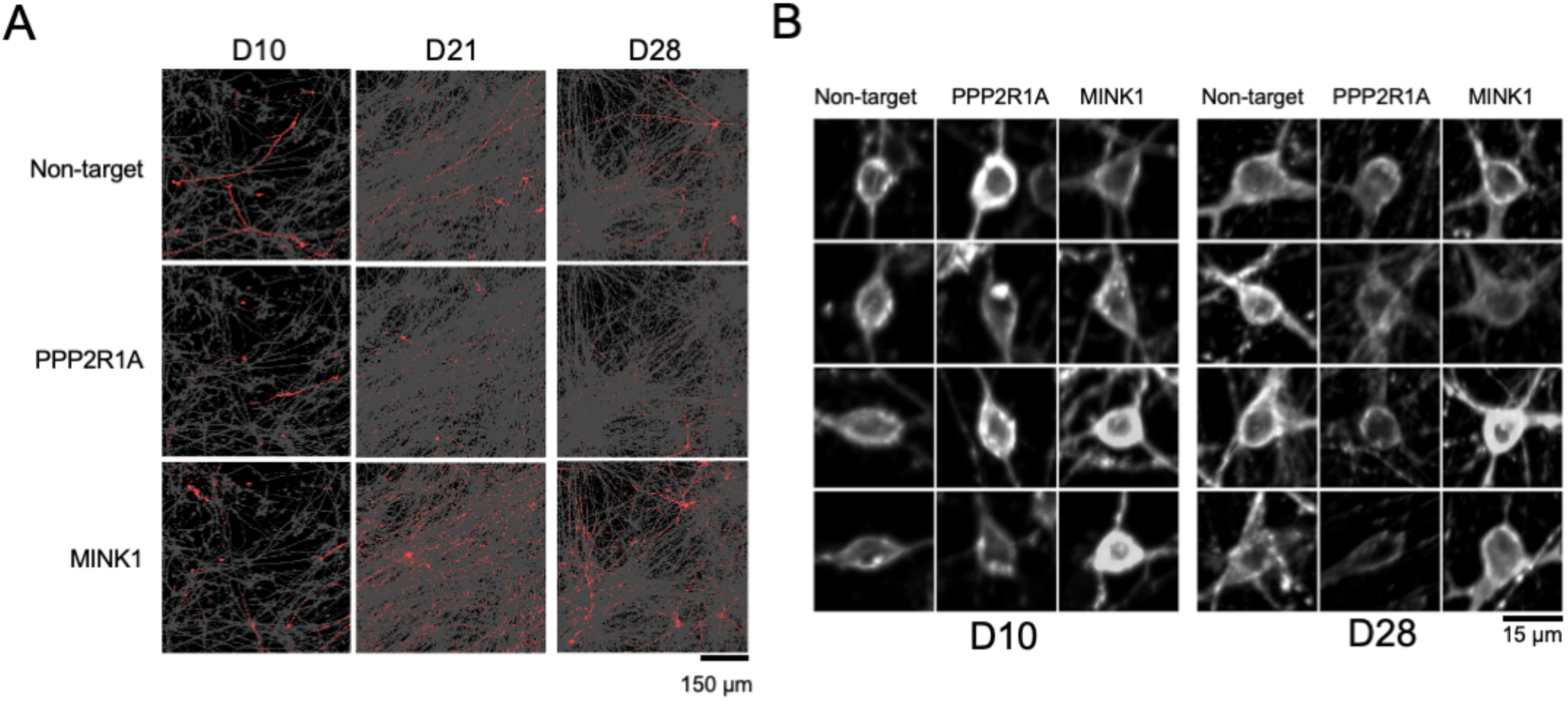
A) Example deconvolved Pro-Code images demonstrating knockdown effects on neurite length and branches. For each timepoint (columns), single field-of-view image is presented with each genotyped cells (non-target, PPP2R1A, MINK1) in red (rows), and all other genotypes in gray. B) Example single-cell cropped images demonstrating effects of knockdown of genes (columns) on neurite trunks.

## Notes

### Competing Interest Statement

The authors have declared no competing interest.

### Summary of Updates

Method name is changed from CellFIE to CellPHIE

